# Systematic Discovery of Motif-based Interactions of the Auxiliary Domains of USP Family Deubiquitinases

**DOI:** 10.1101/2025.09.22.676098

**Authors:** Aimiliani Konstantinou, Alicia Córdova-Pérez, Julia K. Varga, Priyanka Madhu, Leandro Simonetti, Maximilian Vieler, Ryosuke Ishimura, Frederic Lamoliatte, Ora Schueler-Furman, Norman E. Davey, Yogesh Kulathu, Ylva Ivarsson

## Abstract

The ubiquitin-specific proteases (USPs) family is the largest family of human deubiquitinating enzymes (DUBs). While most USPs are agnostic to polyubiquitin linkage-type, their substrate specificity is thought to be mediated by the recognition of the ubiquitylated protein itself. In addition to their catalytic domain, USPs have one or more auxiliary domains (ADs) with key functions in regulating DUB activity and subcellular localization. We hypothesize that some ADs bind short linear motifs (SLiMs) typically found in intrinsically disordered regions of proteins to achieve targeting to substrates and multiprotein complexes. To test this hypothesis, we systematically assessed the potential of 29 USP-ADs and two full-length USPs for SLiM binding using a combination of proteomic-peptide phage display, peptide arrays and affinity measurements. We discovered SLiM-based interactions for 14 ADs from 9 USP-DUBs, including CYLD, USP11, USP19, USP20, USP22 and USP33, and define the consensus motif and properties of the SLiM-AD binding. Interestingly, we established that the zf-UBP and DUSP2 domains of USP20 and USP33 are SLiM binding ADs with similar binding profiles, explaining the functional redundancy between the two DUBs. Our work reveals unique motifs recognized by the auxiliary domains CAP-Gly, UBL, zf-UBP and DUSP, with potential functional implications for substrate recognition and complex assemblies.

## Introduction

Approximately 100 human deubiquitinating enzymes (DUBs) contribute to controlling the ubiquitination status of the cell^1^. They are responsible for removing ubiquitin from a vast number of ubiquitination sites by cleaving either the covalent bond between ubiquitin and the substrate or within polyubiquitin chains. Substrate specificity of DUBs is often achieved by recognition of specific ubiquitin linkage types or ubiquitin chain length^2,3^. However, many ubiquitin-specific proteases (USPs) demonstrate promiscuity in recognising linkage types, rather demonstrating specificity for the substrate that is ubiquitylated.

With 58 members, the USP family represents the largest human family of DUBs. Most USPs possess a modular architecture featuring auxiliary domains (ADs) in addition to their catalytic core that enable substrate recognition independently of the linkage type. Some of the ADs have been shown to bind to short linear motifs (SLiMs), which are typically 3-10 amino acid stretches located within intrinsically disordered regions (IDRs)^4,5^. These interactions may contribute to substrate targeting or complex assembly. A well-known example is USP7, which has an N-terminal MATH (meprin and TRAF homology) domain and a UBL (ubiquitin-like) domain that bind to distinct SLiMs ([PAE]xxS and KxxxK, respectively)^6–8^. Another example is USP8, which features an N-terminal MIT (Microtubule-Interacting and Trafficking) domain, which binds to an MIT interacting motif 1-like SLiM ([DE][LIF]x{2,3}R[FYIL]xxL[LV]) found in ESCRT-III complex proteins and targets the protein to endosomes^9,10^. USP8 also has a catalytically inactive rhodanese domain recently found to engage in SLiM-based interactions with potential substrates^11^. Other USPs with reported SLiM-binding ADs include the cylindromatosis protein (CYLD), which has three cytoskeleton-associated protein-glycine-rich (CAP-Gly) domains of which one reportedly binds to a proline-rich peptide in NEMO (NF-kappa-B essential modulator; also called IKK-gamma)^12^ and USP11 which has a SLiM-binding tandem DUSP (domain present in ubiquitin-specific proteases)-UBL domain^13^. Additionally, the zf-UBP (zinc finger-ubiquitin binding protein) domains of USP5 and USP16 bind to the C-terminal diglycine motif of ubiquitin^14,15^. Despite these emerging insights, the SLiM-binding potential of ADs of most human DUBs remains largely uncharacterized. Uncovering these interactions could provide a deeper understanding of how DUBs achieve substrate specificity, coordinate with cellular complexes, and regulate diverse signalling pathways through motif-guided recognition.

In this study, we systematically screened for SLiM-binding of 29 ADs from 19 USPs and two full-length USPs (Suppl. Table 1). Proteins were screened against a peptide-phage library that tiles the intrinsically disordered regions of the human proteome (called HD2)^16^. We identified potential peptide ligands for 15 of the protein baits. For 11 of the domains, we defined consensus binding motifs through an integrated approach combining ProP-PD data, structural modelling, and peptide SPOT array alanine scanning. Notably, for USP20 and USP33, we discovered that both their zf-UBP and DUSP2 domains act as peptide-binding modules, each capable of interacting with a wide range of peptides. Moreover, many of the interacting proteins and potential substrates contain multiple candidate SLiM-binding sites, suggesting a putative cooperative mode of interaction. Collectively, our findings uncover previously unrecognized peptide-binding properties across several USP auxiliary domains and point to broader functional and regulatory roles for SLiM-mediated interactions in the context of deubiquitination.

## Results and discussion

### Identification of peptide ligands of auxiliary domains of USPs

We generated a collection of 29 USP ADs and two full-length USPs (CYLD and USP7). The collection was biased towards domains that are present in several USPs, such as zf-UBP domains, UBL domains and DUSP domains (Suppl. Table 1). We also included ADs domains only found in specific USPs, such as the three CAP-Gly domains of CYLD and the C-terminal domain of USP25. The proteins were used as baits in ProP-PD selections against the previously described HD2 library (Fig. 1A), which displays overlapping 16-amino-acid peptides tiling the intrinsically disordered regions of the human proteome ^16^. The peptide-coding regions of the phage-pools were analyzed by next-generation sequencing (NGS), mapped to the corresponding proteins and filtered for high/medium confidence ligands based on established criteria (with confidence level 4 being the highest). In total, we found 502 peptides binding to 14 bait protein domains and the full-length proteins CYLD and USP7 (Fig. 1B, C, Suppl. Table 2).

**Figure 1.**
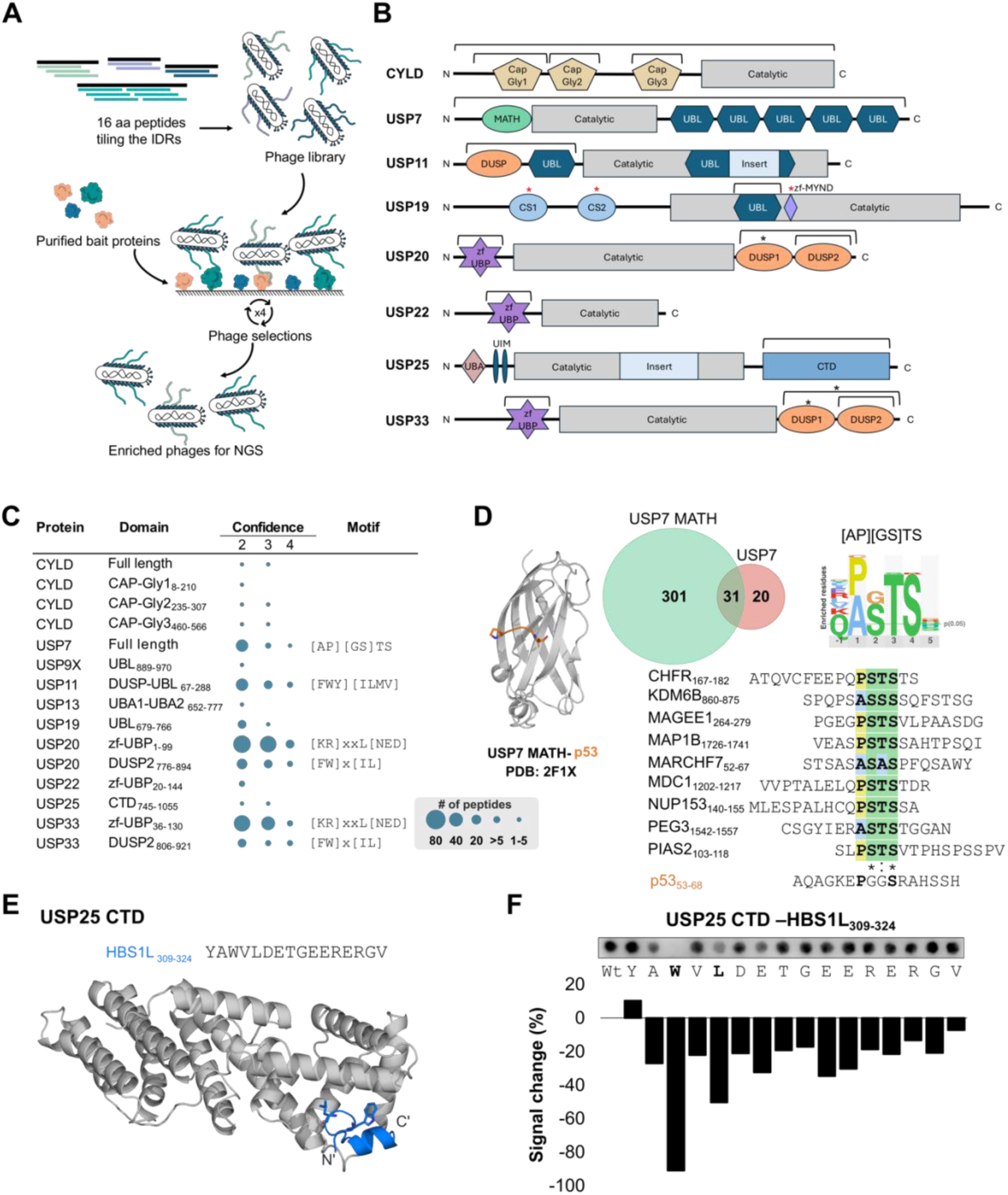
Overview of the ProP-PD selection results for auxiliary domains of DUBs and the full-length CYLD and USP7, including examples of DUBs binding to peptides and motifs. **A.** Overview of the ProP-PD selection. **B.** Schematic representation of the domain composition of the DUBs investigated. The protein or protein domains that were successfully produced and used as baits in phage selections are indicated by square brackets. Domains that were screened without enrichment of binding phages are marked with a red asterisk. Domains that could not be purified are marked with a black asterisk. **C.** Overview of the ProP-PD selections results. The number of peptides enriched for each bait is proportional to the area of the circle. The confidence levels of the peptides identified (with 4 being the highest quality data), and the regular expression of the consensus binding motifs established are indicated. **D.** Peptide binding of USP7. Left: Structure of the USP7 MATH domain binding to the PxxS containing p53 peptide (PDB: 2F1X^6^). Middle: Overlap between previous ProP-PD data for the USP7 MATH domain^16^ and the results obtained for the full-length USP7 in the current study. Right: Position-specific scoring matrix (PSSM) and representative USP7 binding peptides found in this study (the asterisk indicates full conservation and the semicolon semi-conserved residues). The p53 peptide is shown for comparison. **E.** Peptide binding of the C-terminal domain of USP25. AF3^17^ model of USP25 CTD binding to a peptide from HSB1L_309-324_. The WxL motif identified by the alanine scanning (see panel **F**) is docked into a pocket of the domain (ipTM: 0.7). **F.** Alanine scanning SPOT array of the HSB1L_309-324_ peptide.

Analysing the peptides using the SLiMFinder algorithm^18^ revealed six consensus motifs (Fig. 1C). Among them, full-length USP7 preferentially binds to a [AP][GS]TS motif. This motif matches the consensus USP7 MATH domain binding motif, found for example in the tumour suppressor p53, a bona fide substrate of USP7^6^ (Fig. 1D). We previously used the USP7 MATH domain as bait in ProP-PD selections and validated that it binds to a [AP][GS]xS motif ^16^. Here, we found 51 ligands, of which more than 60 % overlap with those identified for the isolated USP7 MATH domain (Fig. 1D). The identified peptide ligands were consistently modelled to bind to the MATH domain binding pocket, when using either the isolated MATH domain or the full-length USP7 for AlphaFold3 (AF3) modelling^17^ (Suppl. Fig. 1A, B).x2 Our screen unravelled binding sites of USP7 MATH domain in several previously reported USP7 interactors, such as the lysine-specific demethylase 6B (KDM6B), the mediator of DNA damage checkpoint protein 1 (MDC1), and the E3 ubiquitin-protein ligases CHFR and MARCHF7^19–22^ (Fig. 1D). CHFR and MDC1 are also previously validated USP7 substrates^22,23^. The USP7 results highlight the robustness of the ProP-PD approach in identifying relevant interactions.

For eight of the ADs, we identified limited sets of peptide ligands and no clear consensus motifs (Fig. 1C). Nevertheless, these hits provided valuable starting points for motif discovery. For example, the C-terminal domain (CTD) of USP25 enriched a peptide region from the HBS1-like protein (HBS1L) spanning residues 305–324 (ASFA<u>YAWVLDETGEER</u>ERGV, with the underlined stretch appearing in two overlapping peptides). An AF3^17^ model of the USP25 CTD–HBS1L interaction (ipTM: 0.7) suggests that the WxL motif docks into a pocket formed by the N-terminal helices of the domain (Fig. 1E, Suppl. Fig. 1C). This interaction was further validated using a peptide alanine-scanning peptide array, which confirmed the importance of the WxL motif for binding (Fig. 1F). These findings illustrate how even a limited number of peptide hits can be mined to reveal previously unrecognized, motif-driven interactions.

### SLiM-based interactions of the CYLD CAP-Gly domains

CYLD is a tumor suppressor that regulates the stability of NF-κB and numerous other targets^24^. It has three CAP-Gly domains, a domain type which is known to interact with peptide ligands and typically recognize an acidic C-terminal motif defined as [ED]x{0,2}[EDQ]x{0,1} [YF]-COOH^25^ (Suppl. Fig. 2A). In CYLD, the CAP-Gly1 and CAP-Gly2 domains facilitate interaction with microtubules, while the CAP-Gly3 domain is known to bind a peptide from NEMO (also called IKKgamma)^12^. In addition, the CAP-Gly2 and CAP-Gly3 domains have been reported to bind to ubiquitin (Suppl. Fig. 2D)^26^. To identify peptide ligands binding to CYLD, we screened the individual CAP-GLY domains through ProP-PD and found one to four ligands per domain (Fig. 2A-C). We confirmed the top hits of each domain using alanine scanning peptide arrays. The results validated the interaction of the CAP-Gly1 domain with a peptide from the RNA-binding protein 25 (RBM25), and showed it to bind via an atypical CAP-Gly binding motif: LxxMxxxAxxRR (Fig. 2A). AF3 modelling (ipTM: 0.93) predicted that the peptide adopts an α-helical conformation upon binding the first CAP-Gly1 domain of CYLD (Fig. 2A). The CAP-Gly2 domain was confirmed to bind to a peptide from FBXO46 (F-box only protein 46; 168-DLLSVAEMVALVEQRA-183), and this interaction was found to be sensitive to mutations at most peptide positions (Fig. 2B). The interaction between CYLD CAP-Gly2 and the FBXO46 peptide was modelled with high confidence using AF3 (ipTM: 0.84) (Fig. 2B). Notably, similar to the CAP-Gly1 binding peptide, the CAP-Gly2 binding peptide was modelled in alpha-helical conformations, but oriented in the opposite direction (Fig. 2A, B). The CAP-Gly3 domain binds a peptide from the probable E3 ubiquitin-protein ligase HECTD4. Alanine scanning peptide array showed that a GxFKDEIYIP stretch was critical for binding (Fig. 2C). We modelled the CYLD CAP-Gly domains interactions using AF3 (Fig. 2A-C), resulting in high-confidence models (ipTM = 0.77 - 0.93). The HECTD4-derived peptide was confidently modelled binding to the CAP-Gly3 domain in a β-hairpin conformation (ipTM: 0.77; Fig. 2C). The binding site of the HECTD4 peptide on the domain partially overlaps with the peptide binding site suggested by NMR for the NEMO peptide^12^. The results suggest a conserved binding surface with potential adaptability depending on the peptide ligands.

**Figure 2.**
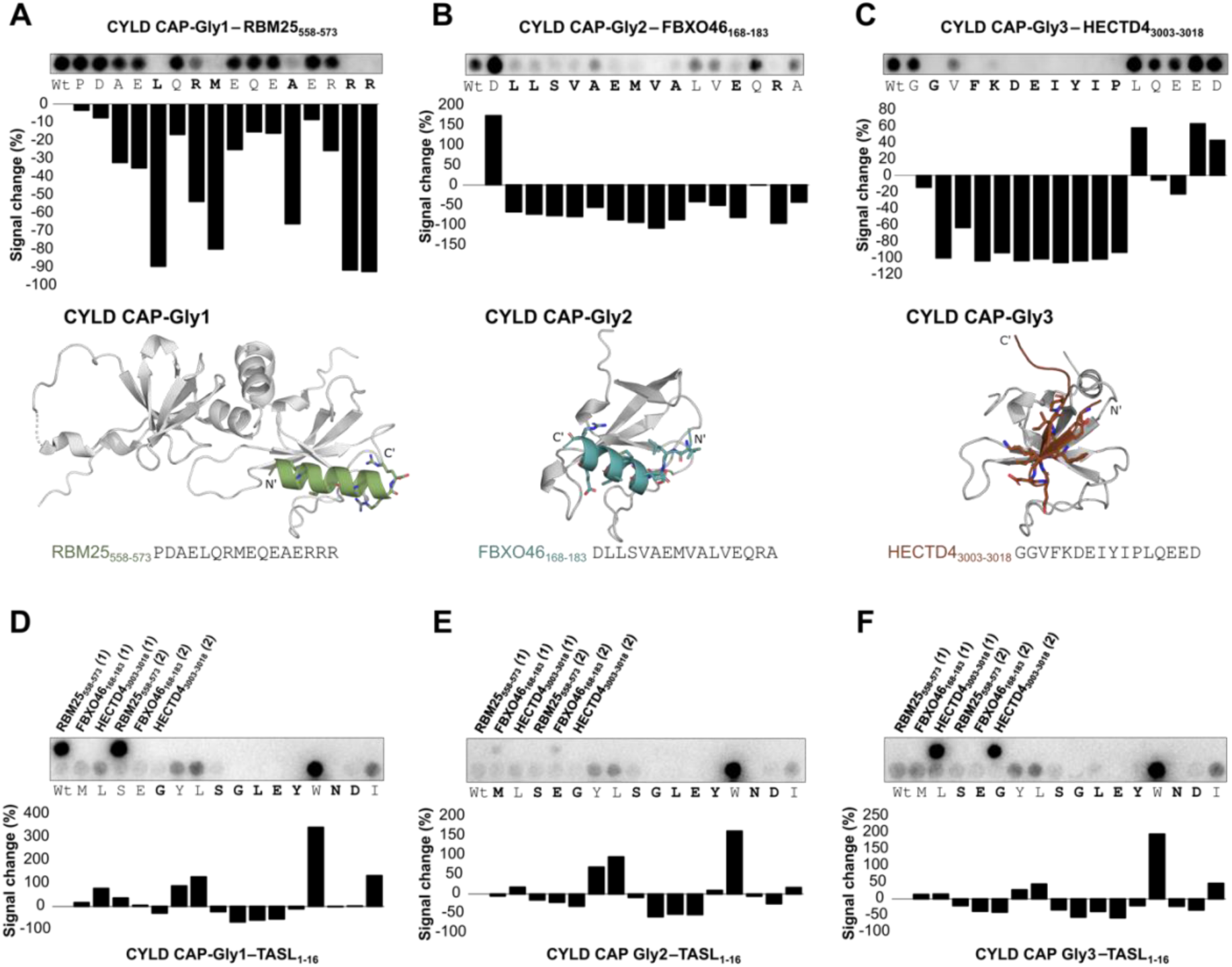
The three CAP-Gly domains of CYLD are peptide binding domains. **A.** Peptide binding of CYLD CAP-Gly1. Alanine scanning peptide array (top) confirms an interaction between CYLD CAP-Gly1 domain and a RBM25_558-573_ peptide and identifies residues critical for binding. The AF3 model of the complex (bottom; ipTM: 0.93) illustrates that the peptide forms an α-helix upon binding and supports that the identified residues are making contacts with the domain. **B.** Peptide binding of the CAP-Gly2 domain of CYLD. Alanine scanning peptide array (top) confirms an interaction between CYLD CAP-Gly2 and an FBXO46_168-183_ peptide. The AF3 model of the complex (bottom; ipTM: 0.84) suggests that the peptide forms an α-helix upon binding. **C.** Peptide binding of the CAP-Gly3 domain of CYLD. Alanine scanning peptide SPOT array (top) confirms an interaction between CYLD CAP-Gly3 and an HECTD4_3003-3018_ peptide, and that the interaction is sensitive to mutations of an extended stretch. The AF3 model of the complex (bottom; ipTM: 0.77) illustrates that the peptide forms a β-hairpin structure upon binding. **D-F.** Alanine scanning peptide array analysis of the TASL peptide binding to the CYLD CAP-Gly1 (**D**), CAP-Gly2 (**E**) and CAP-Gly3 (**F**) domains. **D-F.** Peptides from panel A-C used as positive and negative controls for each domain.

To get additional information, we screened the full-length CYLD for peptide binding, which resulted in a set of 10 peptides. Of these, two highly enriched and overlapping peptides belong to TASL (TLR adapter interacting with SLC15A4 on the lysosome). A peptide array analysis revealed that all CAP-Gly domains bind to the TASL peptide (Fig. 2D-F, Suppl. Fig. 2E-G). Thus, the selection against the CYLD full-length protein enriched a peptide that can bind all the three CAP-Gly domains. For the CAP-Gly2 domain, the spot intensity was higher for the TASL peptide as compared to the FBXO46 peptide, potentially suggesting a higher affinity for the TASL peptide. On the contrary, the RBM25_558-573_ and HECTD4_3003-3018_ peptides appeared to be better binders of the CAP-Gly 1 and 3 domains, respectively. Alanine scanning revealed that the key residues for TASL peptide binding to the three domains converge on a GLEYxND stretch (Fig. 2D-F). Notably, a tryptophan to alanine (W13A) mutation of the wild-card position enhanced the binding to all three domains. Taken together, we find that all three CYLD CAP-Gly domains are peptide-binding ADs, being bound by distinct, as well as shared peptide ligands.

### SLiM-based interactions of the ubiquitin-like (UBL) domains of USP19 and USP11

Ubiquitin-like (UBL) domains are homologous to ubiquitin and found in one third of the USP DUBs. UBL domains are known to bind the ubiquitin interacting motif (UIM; [DE]x{2,4}[AV]xx[LMIV]xx) that forms an α-helix upon binding^27,28^. To explore novel interactions mediated by UBL domains of the USP DUBs, we screened five UBL domains and identified binders for the UBL domain of USP19 and the tandem DUSP-UBL domain of USP11. For the USP19 UBL domain, the most enriched peptide was from the protein Hook homolog 2 (HOOK2) (Suppl. Table 2; Fig. 3A), a sequence that is conserved in the homologous protein, HOOK1. Peptide array alanine scanning analysis of a HOOK2 peptide (590-DADLRAMEERYRRYVD-605) and a peptide from the protein KIBRA (WWC1; 1029-ELPQWLREDERFRLLL-1044) identified partially shared binding determinants (LRx{3,4}R[FY]) (Fig. 3B) that are distinct from the typical UIM motif. AF3 modelling of the HOOK2 and WWC1 peptides with the USP19 UBL domain revealed that the peptides adopt α-helical conformations (Fig. 3C). Notably, the binding sites of the model peptides partially overlap with the UIM binding site of ubiquitin, although the sequence identities between the domains are low (Suppl. Fig. 3A) and the I44 in the central β-strand of the ubiquitin binding site, which is important for UIM binding^29^, is replaced by a valine in the USP19 UBL domain.

**Figure 3.**
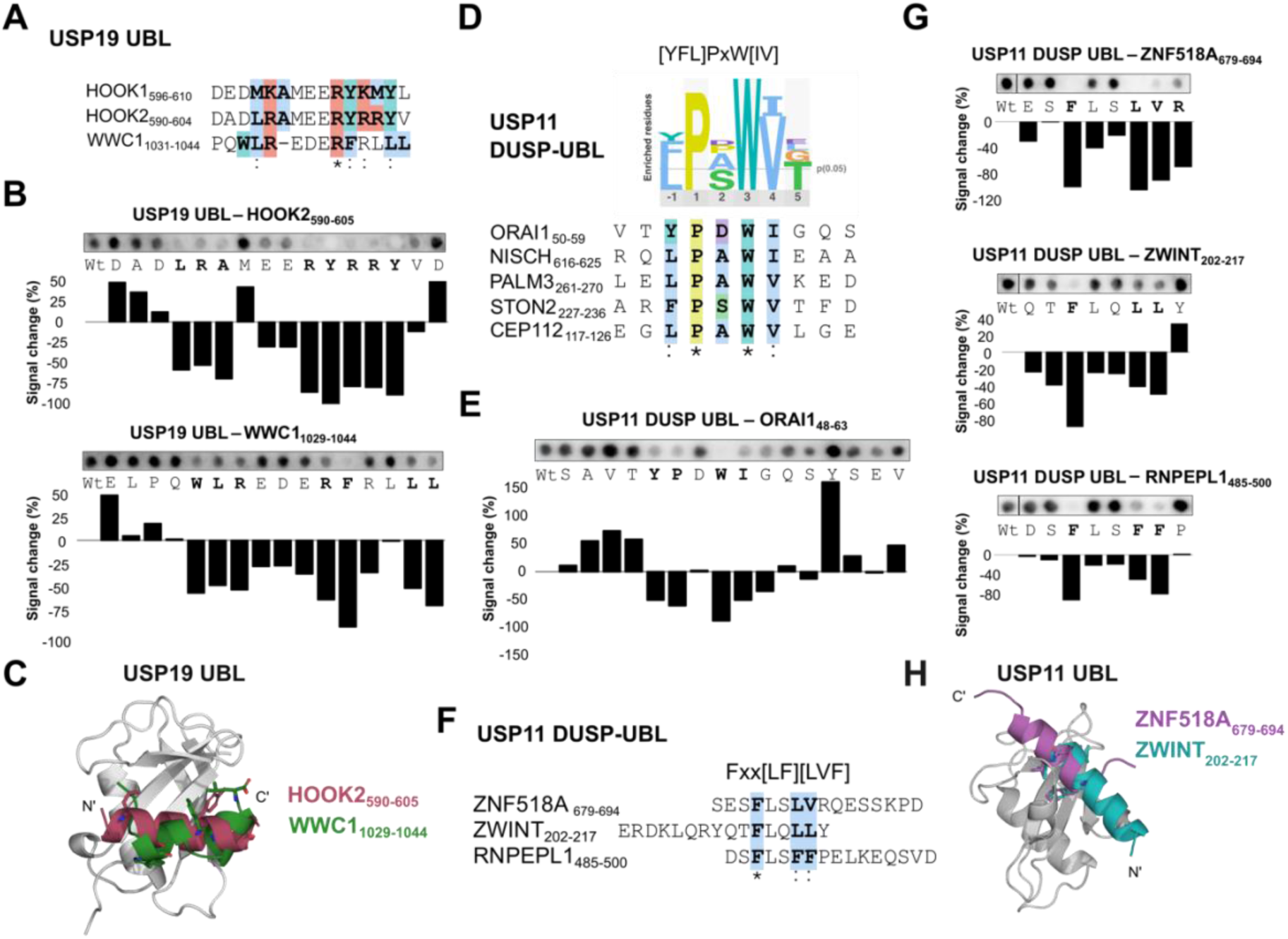
**A.** Alignment of the USP19 UBL binding peptides from HOOK2 and WWC1 identified through ProP-PD selections, together with the conserved region from the HOOK2 homologous protein HOOK1. The asterisk indicates full conservation and the semicolon semi-conserved residues. Additional USP19 UBL peptides are available in Suppl. Table 2. **B.** Alanine scanning peptide array analysis of two USP19 UBL binding peptides from HOOK2_590-605_ and WWC1_1029-1044_. **C.** Overlay of the AF3 models of USP19 UBL domain and the HOOK2_590-605_ and WWC1_1029-1044_ peptides (ipTMs: 0.78). Key residues identified by alanine scanning are shown in stick representation. **D.** Alignment of five USP11 DUSP-UBL domain binding peptides with an alternative [YFL]PxW[IV] motif, together with the generated PSSM. The asterisk indicates full conservation and the semicolon semi-conserved residues. **E.** Alanine scanning peptide SPOT array of the interaction between USP11 DUSP-UBL and the ORAI1_48-63_ peptide. **F.** Alignment of three USP11 UBL binding peptides from ZNF518A, ZWINT and RNPEPL1. Additional USP11 DUSP UBL binding peptides are available in Suppl. Table 2. The asterisk indicates full conservation and the semicolon semi-conserved residues. **G.** Alanine scanning peptide SPOT array of the USP11 DUSP-UBL domain binding peptides ZNF518A_679-694_, ZWINT_202--217_ and RNPEL1_485-500_. Partial arrays are shown to highlight the identification of a common Fxx[LF][LVF] motif (see Suppl. Fig. 3C for the complete array results). **H.** AF3 models of USP11 DUSP-UBL domain with the ZNF518A_679-694_ (ipTM: 0.71), ZWINT_202--217_ (ipTM: 0.66) peptides. The peptides are predicted to dock to the UBL domain. The motif key residues are shown in stick representation (see Suppl. Fig. 3E for more details).

Conversely, the USP11 DUSP-UBL tandem domain enriched for a diverse set of ligands, 17 of which contained a consensus [YFL]PxW[IV] motif (Fig. 3D). Alanine scanning analysis of a peptide from the calcium release-activated calcium channel protein 1 (ORAI1) (48-SAVTYPDWIGQSYSEV-63) confirmed the YPxWI motif (Fig. 3E). AF3 modelling of the DUSP-UBL ORAI1 complex resulted in low confidence models, likely reflecting a low affinity interaction, as determined by FP-based affinity measurements (K_D_: 200-300 μM; Suppl. Fig. 3B). Additionally, the selection against the USP11 DUSP-UBL tandem domain enriched another cohort of peptides with an apparent [LF]xx[LF][LVF] motif (Fig. 3F), which is similar to a previously characterized USP11 binding motif^13^. Key motif residues were confirmed in peptides from ZNF518A, ZWINT and RNPEPL1 by alanine scanning (Fig. 3G, Suppl. Fig. 3C). AF3 confidently modelled the ZNF518A (ipTM: 0.71) and ZWINT (ipTM: 0.66) peptides to a common site in the USP11 UBL domain (Fig. 3H) which overlaps with a previously established USP11 UBL binding site for a AEGE**F**YK**L**KIRTPQ peptide (PDB: 5OK6; Suppl. Fig. 3D)^13^. The phenylalanine and the first leucine of the motif appear to be oriented towards the binding pocket (Suppl. Fig. 3E). Taken together, we found that the USP11 DUSP-UBL domain binds to two types of SLiMs, the previously described [LF]xx[LF][LVF] motif and the newly identified [YFL]PxW[IV] motif, representing an additional type of USP19 binding motif.

### The zf-UBP domains of USP20 and USP33 recognize a [KR]xxL[NED] motif

USP20 and USP33, also known as VHL-interacting deubiquitinating enzymes hVDU2 and hVDU1, are ER-localized DUBs that regulate various cellular biological processes, such as cell cycle progression and proliferation ^30–32^. USP20 and USP33 have a similar domain organization (Fig. 1B). They consist of four domains: an N-terminal zf-UBP domain, a catalytic domain, and two closely linked ubiquitin specific proteases 1 (DUSP1) and (DUSP2) domains (Fig. 1B). We screened the zf-UBP and the DUSP2 the domains of USP20 and USP33, as well as the DUSP1-DUSP2 tandem of USP20. For the zf-UBP domains we identified 136 and 111 peptide-binding regions, respectively, with an overlap of 67 peptides binding both domains (Fig. 4A).

**Figure 4.**
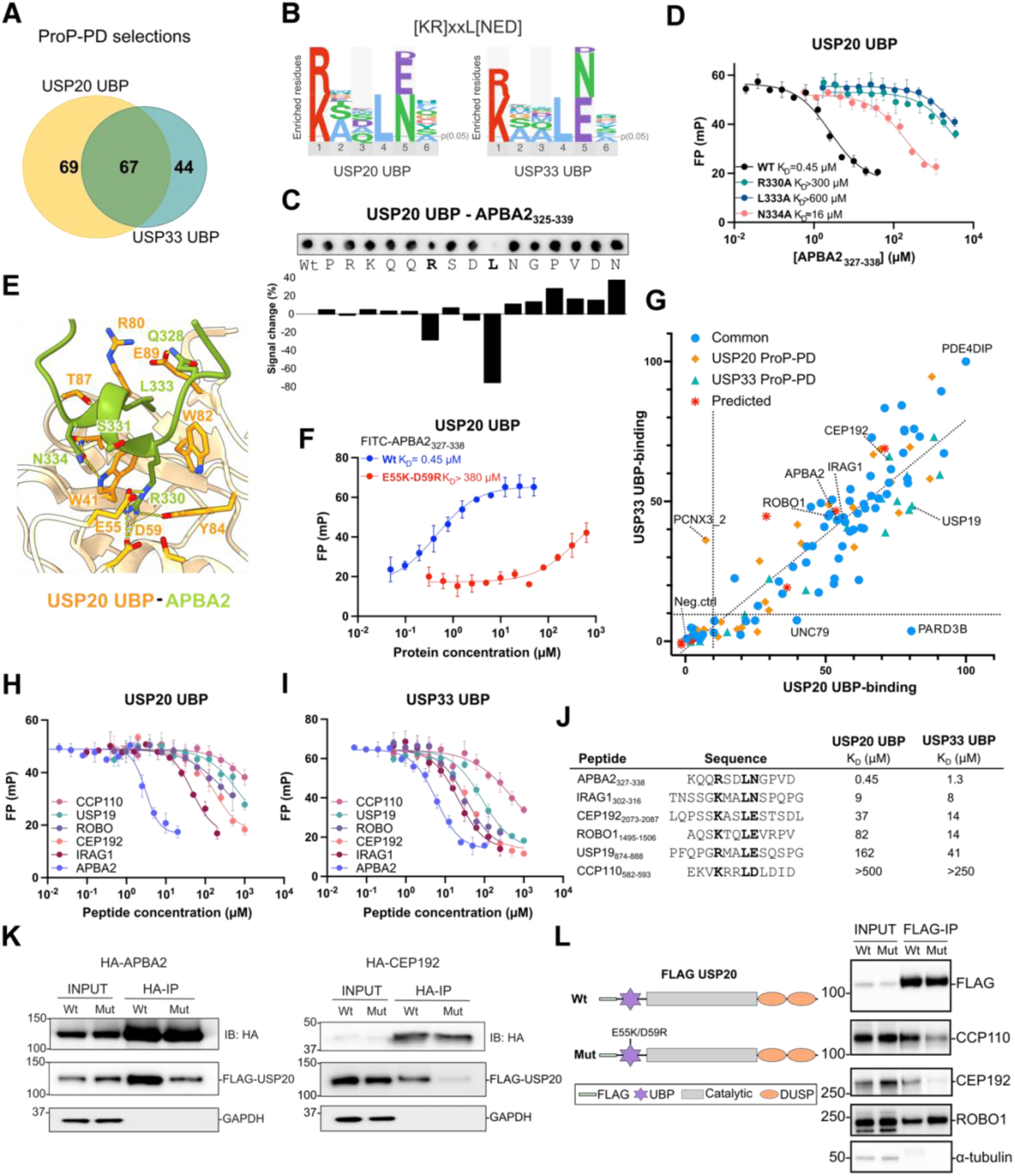
The zf-UBP domains of USP20 and USP33 bind to [KR]xxL[NED] motif containing peptides. **A.** Venn diagram of the overlap of the peptide-binding regions identified as ligands for the zf-UBP domains of USP20 and USP33 through ProP-PD selections. **B.** Consensus binding [KR]xxL[NED] motif for USP20 and USP33 zf-UBP domains. **C.** Peptide array alanine scanning of the APBA2_325-339_ peptide binding to the USP20 zf-UBP domain. **D.** Competitive FP-monitored affinity measurements of USP20 zf-UBP binding to wild-type and mutant APBA2_327-338_ peptides (n = 3). **E**. AF3 model of the interactions between the APBA2_327-338_ peptide and the zf-UBP domain of USP20 (pLDDT> 80 over the motif residues). **F.** FP-monitored affinity measurements between wild-type and double mutant zf-UBP domains of USP20 and the APBA2_327-338_ peptide (n= 3). **G.** Scatter plot of the normalized peptide SPOT intensities observed for 120 peptides against the zf-UBP domains of USP20 and USP33. The peptides used to design the array were found in phage selections against both USP20 and USP33 (“Common”), in either of the two USP20 and USP33 zf-UBP domain datasets or were predicted ligands based on the consensus motif. The diagonal line represents the linear regression of the data (slope: 0.82). The dotted lines indicate a cut-off of 10% signal over background. Peptide sequences and peptide array analysis results are available in the Suppl. Table 4. **H-J.** Competitive FP-monitored affinity measurements of USP20 (**H**) and USP33 (**I**) zf-UBP domains binding to six distinct peptides (**J**), together with the estimated K_D_ values (n= 3). **K.** Co-immunoprecipitation experiments of FLAG-USP20 with wild-type HA-APBA2 and mutant (R330A/L333A) and CEP192 K1743-K2092 and mutant (K2078A/L2081A) in HEK293 cells. **L.** Immunoprecipitation of wild-type USP20 and predicted zf-UBP binding pocket mutants. Left: Schematic representation of the USP20 zf-UBP predicted binding pocket mutants (E55K/D59R). Right: Immunoprecipitation experiments of the FLAG-USP20 with endogenous CCP110, CEP192 and ROBO1.

Based on the identified peptides, consensus motifs were generated using the SLiMFinder algorithm^18^ which converged on a novel [KR]xxL[NED] motif (Fig. 4B). Interestingly, for many of the ligands, ubiquitination sites are present in and around the motif (Sup. Table 2). To validate the importance of each residue in this motif to zf-UBP binding, we performed an alanine scanning analysis of a peptide (325-PRKQQ**R**SD**LN**GPVDN-339) from amyloid-beta A4 precursor protein-binding family A member 2 (APBA2; also known as MINT2), a previously reported interactor of USP20/USP33 identified in affinity-purification mass spectrometry (AP-MS) and yeast two-hybrid (Y2H) experiments^33–35^. The analysis confirmed binding and established a key requirement for leucine at the 4^th^ position of the motif (Fig. 4C). Furthermore, an R to A mutation of the first position of the motif affected the interaction. To further assess the motif-determinants we conducted FP-monitored affinity measurements of wild-type and mutant APBA2_327-338_ peptides against the zf-UBP domains of USP20 and USP33 (Fig. 4D; Supp. Fig. 4A). While the wild type APBA2 peptide bound with high affinity (USP20 K_D_= 0.45 μM; USP33; K_D_=1.3 μM), mutation of the arginine at the first position of the motif (R330A; USP20: K_D_> 300 μM, USP33: K_D_>500 μM) and the leucine at position 4 (L333A; USP20: K_D_> 600 μM, USP33: K_D_=ND) significantly weakened binding (Fig. 4D, Supp. Fig. 4A). Additionally, the asparagine at position 5 contributes to binding, as revealed by a N334A mutant conferring reduced affinity for USP20 and USP33, albeit to less extent (USP20 K_D_≈ 16 μM; USP33 K_D_≈ 49 μM) (Fig. 4D; Supp. Fig. 4A). Taken together, the USP20/33 zf-UBP interaction is centred on the [KR]xxL[NED] motif.

AF3 docking of the APBA2_327-338_ peptide, with USP20/USP33 UBP domains resulted in high confidence models (Suppl. Fig. 4B; pLDDT> 80 over the motif residues). The [KR]xxL[NED] peptide binding pocket of these domains overlaps with the binding site for C-terminal ubiquitin present in other zf-UBP domains, such as USP5 and USP16^15,36^ (Suppl. Fig. 4C). In contrast to most other zf-UBP domains, the USP20 and USP33 zf-UBP domains do not bind the ubiquitin C-terminus with measurable affinity (Suppl. Fig. 4D). Nevertheless, the interactions between the APBA2_327-338_ peptide and the USP20/USP33 zf-UBP domain (Fig. 4E, Suppl. Fig. 4E), have some resemblance to how the USP5 zf-UBP binds to the ubiquitin C-terminus (PDB: 2G45)^15^. Residues W41, W82, Y84 in USP20 (W72, W113, Y115 in USP33) form a hydrophobic pocket similar to the one formed by W209, Y259 and Y261 in USP5 (Suppl. Fig. 4C). Moreover, USP20 E55 (E86 of USP33) interacts with the arginine at the first position of the [KR]xxL[NED motif and forms an additional salt bridge with USP20 D59 (D90 in USP33). To validate the peptide binding site, we created double mutants of the USP20 and USP33 zf-UBP domains (E55K/D59R and E86K/D90R, respectively), and affinity measurements confirmed that the mutations significantly impair peptide binding (Fig. 4F, Suppl. Fig. 4E).

To assess the binding of a larger set of USP20/USP33 zf-UBP ligands, we designed an array that included peptides that bind to both USP20 and USP33 zf-UBP domains based on the phage selection results (Fig. 4G, Suppl. Table 3E). In addition, the array design included “unique” peptides only found as ligands for either of the zf-UBP domains. An additional set of nine predicted ligands (based on PSSMSearch^37^) were included. More than 80% of the peptides were found to bind to USP20, while over 70% of the peptides bind to the USP33 zf-UBP, with a signal intensity exceeding 10% relative to the background (Fig. 4G; Suppl. Fig. 4G). Applying a more stringent cut-off (20% signal over background), 79 ligands were confirmed to bind to one or either domain. Four of the nine predicted ligands were confirmed to bind the zf-UBP domains, suggesting that USP20/33 zf-UBP binding peptides can be predicted using the motif determinants with some accuracy. Importantly, most of the peptides bound to both domains, with a peptide from the partitioning defective 3 homolog B protein (PARD3B) (171-TQNLEDREVLNGVQTE-186) being an exception, as it only bound to USP20.

Amongst the validated ligands we note a peptide from Roundabout homolog 1 (ROBO1_1495-1506_; see Suppl. Fig. 4F for AF3 model). ROBO1 is known to be deubiquitinated by USP33 in the Slit-ROBO signalling pathway and regulates cell migration^38,39^. The region of interaction between ROBO1 and USP33 has previously been mapped to the C-terminal stretch of ROBO1, which overlaps with the peptide discovered here. We found that the ROBO1_1495-1506_ peptide binds with almost six-fold higher affinity to USP33 zf-UBP over the USP20 zf-UBP domain, with K_D_ values of 14 μM and 82 μM, respectively (Fig. 4H-J). We further determined the affinities for a set of ProP-PD derived ligands and one additional predicted binding peptide from the centriolar coiled-coil protein of 110 kDa (CCP110) (Fig. 4H-J). CCP110 is involved in centrosome duplication and is deubiquitinated and stabilized by USP33 during the S and G2/M phases of the cell cycle ^40^. The affinities were in the range of low- to high-micromolar, with the affinity ranking of the peptides generally being the same for both domains. While the APBA2 peptide binds with the highest affinity (USP20: K_D_=0.45 μM, USP33: K_D_=1.3 μM) to both USP20 and USP33 zf-UBP domains, the predicted CCP110_582-593_ peptide was found to bind with low affinity, again illustrating that while motif-predictions may identify binding sites in known ligands, there is a certain degree of false positives.

To assess the binding of the [KR]xxL[NE] motif in the context of full-length proteins, we co-expressed FLAG-tagged USP20 with either wild-type or motif-mutant HA-tagged APBA2 and CEP192 (the centrosomal protein of 192 kD, amino acid region 1743-2092) and conducted co-IP experiments, which validated interactions with USP20 (Fig. 4K). The interactions were significantly reduced when the [KR]xxL[NE] motif was mutated, as seen for the R330A/L333A mutant HA-APBA2 and the K2078A/L2081A mutant HA-CEP192_1743-2092_ (Fig. 4K). We further generated binding pocket mutants of the zf-UBP in the context of full length USP20 and assessed the effects on binding of FLAG-USP20 to endogenous CCP110, CEP192 and ROBO1 through immunoprecipitation experiments (Fig. 4L). As expected, mutations in the zf-UBP domain of USP20 reduced its interaction with CCP110 and CEP192. However, the mutations did not affect its interaction with ROBO1, which may suggest that the interaction involves additional binding determinants.

Taken together, we established that the zf-UBP domains of USP20 and USP33 are peptide binding modules with largely overlapping specificities for the [KR]xxL[NED] motif-containing ligands, and that the motif is of importance for the interactions between the full-length proteins.

### The DUSP2 domains of USP20 and USP33 bind a [FW]x[IL] consensus motif

The two consecutive C-terminal DUSP domains (DUSP1 and DUSP2) of USP20 and USP33 have been suggested to play a role in substrate recognition^41^. ProP-PD screens of the DUSP2 domain of USP20 and USP33 and the DUSP1-DUSP2 domains of USP20, identified 36 and 38 peptide binders respectively, with 21 common ligands (Fig. 5A; Suppl. Fig. 5A). For the USP33 DUSP2 domain, we identified 34 ligands, of which 18 also bind to the USP20 DUSP2 domain (Fig. 5A). The DUSP2 binding peptides share a [FW]x[IL] motif (Fig. 5B). To validate the consensus motif, we focused on a peptide from the hypoxia-inducible factor 1-alpha (HIF1A), which is a previously reported USP20 substrate^42^. Peptide array alanine scanning confirmed the requirement for a F572 and L574 in the DUSP2 [FW]x[IL] binding motif (Fig. 5C). The importance of the motif residues was further assessed by FP-based affinity measurements, which showed that the HIF1A peptide binds to the DUSP2 domains of USP20 (K_D_=19 μM) and USP33 (K_D_= 1.5 μM) (Fig 5D; Suppl. Fig. 5B) and that mutation of motif residues significantly impacted DUSP2 binding (F572A: USP20 K_D_> 600 μM; USP33 K_D_> 900 μM; L574A: USP20 K_D_= 89 μM; USP33 K_D_> 200 μM). We noticed that the HIF1A peptide contains an acidic stretch of residues immediately upstream of the [FW]x[IL] motif, from D571 (p-1 position relative to motif) to D569 (p-3). A charge-reverting point mutation (D571K) reduced the affinity for the DUSP2 domains, albeit not to the extent as the core hydrophobic residues (D571K: USP20 K_D_= 85 μM; USP33 K_D_= 145 μM) (Fig. 4D). In contrast, a D570K had relatively minor effects on binding. We further noticed that a proline (P564), located just upstream of the DUSP2 binding region in HIF1A is hydroxylated under normoxia. HIF1A proline hydroxylation results in an interaction with the von Hippel-Lindau disease tumor suppressor (pVHL) which subsequently triggers the ubiquitination and degradation of HIF1A^43^. As USP20 counteracts the ubiquitination of HIF1A and stabilizes the protein^42^ we evaluated the effect of hydroxylation on DUSP2 binding but found it to have minor effects on affinities (USP20 K_D_= 25 μM; USP33 K_D_= 7 μM; Fig. 5H-J).

**Figure 5.**
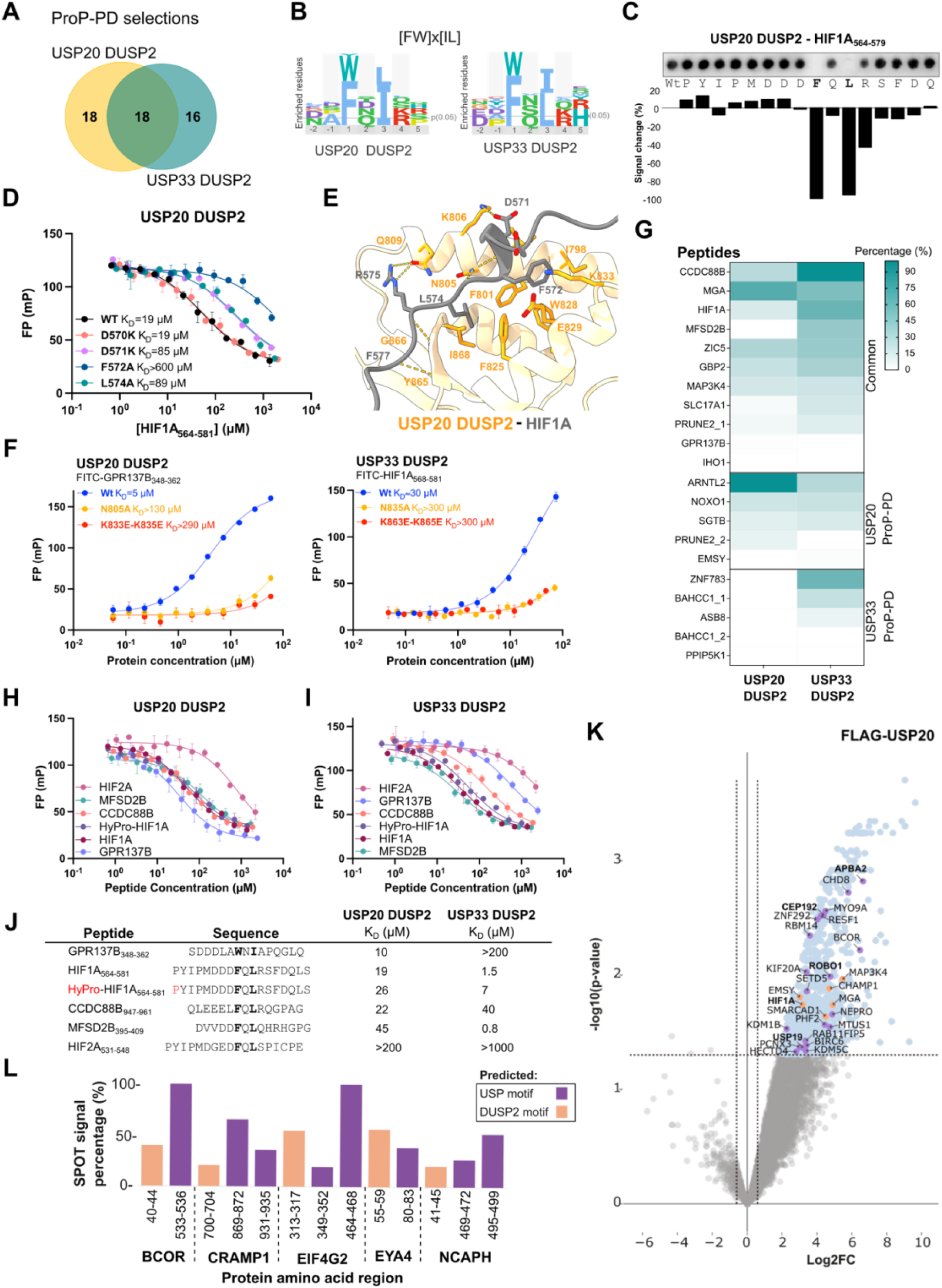
The DUSP2 domains of USP20 and USP33 bind [FW]x[IL] motif containing peptides. **A.** Venn diagram of the overlap of USP20 and USP33 DUSP2 binding peptide-binding regions identified through ProP-PD selections. **B.** Consensus binding motifs for USP20 and USP33 DUSP2 domains generated based on the peptide ligands identified through ProP-PD selections. **C.** Peptide SPOT array alanine scanning of the HIF1A_564-579_ peptide binding to the USP20 DUSP2 domain. **D.** Competitive FP-monitored affinity measurements of USP20 DUSP2 binding to wild-type and mutant HIF1A_564-581_ peptides (n = 3). **E.** AF3 model of USP20 domain binding to HIF1A_564-579_ (pLDDT>90 values over the residues). **F.** Competitive FP-monitored affinity measurements of wild-type, single and double mutant USP20 and USP33 DUSP2 domains binding to FITC-labeled [FW]x[IL] containing probe peptides (n = 3). **G**. Heatmap representation of peptide array analysis of 21 peptides binding to USP20 and USP33 DUSP2 domains. Peptides were found in phage selections against both USP20 and USP33 (“Common”), or in either of the two USP20 and USP33 DUSP2 domain datasets. Peptide sequences and peptide array results are available in the Suppl. Table 5. **H-J.** FP-monitored competitive affinity measurements of USP20 (H) and USP33 (J) DUSP2 domains binding to six peptide ligands. The estimated K_D_ values are provided (n = 3; J). **K.** FLAG-USP20 IP-MS analysis compared to the empty pCMV vector (Suppl. Table 6). Overlap with ProP-PD derived ligands are indicated in purple (zf-UBP) and orange (DUSP2). Interactors validated through FP-monitored affinity measurements are indicated in bold. **L.** Summary of peptide array validation of predicted dual zf-UBP (purple) and DUSP2 motifs (orange) found in IP-MS derived interactors.

AF3 modelling of the HIF1A_564-579_ peptide binding to the DUSP2 domains of USP20 and USP33 revealed the [FW]x[IL] motif (pLDDT>90 values over the residues; Suppl Fig. 5C) to bind in hydrophobic pockets created by F801, I868, F825, E829, W828 in USP20 and F831, I897, F855, E859, W858 in USP33 (Fig. 5E; Suppl. Fig. 5D). Additional electrostatic interactions are mediated between the acidic residues in the HIF1A peptide and K806 and K833 of USP20 (R836 and K863 of USP33). Furthermore, the backbone of the HIF1A peptide is kept in place with H-bonds, partially by complementing a short β-strand of the DUSP2 domain. Similar interactions were observed for a peptide from the integral membrane protein GPR137B (Suppl. Fig. 5E). The DUSP2 peptide binding site was confirmed using single and double mutations of USP20 (N805A and K833E/K835E) and USP33 (N835A, K863E/K865E) (Fig. 5F). We next confirmed the interaction between full-length USP20 with HIF1A (either in normoxia or hypoxia-mimicking conditions) and assessed the effects of mutating the motif (Suppl. Fig. 5F-H). However, the motif mutant (HA-HIF1A F572A-L574A) did not affect binding (Suppl. Fig. 5F). Similarly, mutation of the DUSP2 domain (N804A, K832E, K834E) did not abrogate binding, implying a more stable association (Suppl. Fig. 5G, H). Deletion of either the zf-UBP domain or the DUSP2 domain disrupted recognition of a modified, presumably ubiquitylated, form of HIF1A (Suppl. Fig. 5G, H), suggesting a multivalent mode of interaction.

We designed a peptide array containing 21 peptides to evaluate the binding of a larger set of peptides to the DUSP2 domains. Eleven of the peptides had been identified as ligands for both domains in the phage selection, and five peptides were identified as unique ligands to each domain (Fig. 5G; Suppl. Table 5). Binding was confirmed for 16 out of 21 tested peptides to either domain, with some noticeable specificity differences. In particular, none of the USP33 specific peptides from ProP-PD bind to USP20 DUSP2. In line with this, affinity measurements using an additional set of peptides from the phage selections (GPR137B_348-362_, coiled-coil domain-containing protein 88B (CCDC88B_947-961_), sphingosine-1-phosphate transporter MFSD2B_395-409_) and a predicted DUSP2 binding peptide from endothelial PAS domain-containing protein 1, also known as HIF2A (531-PYIPMDGEDFQLSPICPE-548) (Fig. 5H-J) supported specificity differences between the domains (Fig. 5J). While the UPS20 DUSP2 domain bound most ligands tested with similar affinities, the USP33 DUSP2 domain displayed higher specificity. Also, both domains bound the predicted HIF2A ligand with low affinity.

To substantiate the results, we conducted IP-MS experiments using FLAG-tagged wild-type USP20 and USP20 variants with either the zf-UBP or the DUSP2 domains deleted, and similar experiments for USP33 (Suppl. Table 6). USP20 (and USP33) pulled-down a large number of proteins compared to the negative control (empty vector), including many interactors identified in ProP-PD, such as the APBA2, CEP192, ROBO1, USP19 and HIF1A (Fig. 5K, Suppl. Fig. 5I). Comparison of the results obtained for the full-length and zf-UBP domain mutant variants revealed that the interaction with APBA2 is dependent on the zf-UBP binding pocket (Suppl. Fig. 5J), consistent with the FLAG-IP results of other interactors (Fig. 4L). The deletion of either the zf-UBP or the DUSP2 domains resulted otherwise in surprisingly few changes of the overall interactome profile of USP20 (Suppl. Table 6), suggesting that the interactions of the full-length proteins may involve multiple components (e.g. motif binding of both the zf-UBP domain and the DUSP2 domain, and/or the catalytic domain). We therefore scanned the IP-MS identified USP20 interactors for putative zf-UBP and DUSP2 motifs using the SLiMSearch algorithm and found that 44% of the proteins contain putative binding motifs for both domains in their intrinsically disordered regions (Suppl. Table 7). In contrast, only 27% of the proteins contained neither of the motifs in their IDRs. We tested the binding of 30 and 16 of the predicted binding zf-UBP and DUSP2 SLiMs respectively, using a peptide array analysis, which confirmed 50% of the predicted interactions and the presence of both zf-UBP and DUSP2 binding motifs in several target proteins (Fig. 5L). For example, we validated predicted zf-UBP and DUSP2 binding motifs in the BCL-6 corepressor (BCOR), in the protein phosphatase EYA1 and the eukaryotic translation initiation factor 4 gamma 2 (EIF4G2).

Taken together, we uncover that both the zf-UBP and DUSP2 domains of USP20 and USP33 are SLiM-binding domains, which may contribute to complex assembly and substrate recognition. The results suggest that multiple components may act together to recruit USP20 and USP33 to their substrates. Further investigations are needed to explore the possibility of a cooperative substrate recognition mechanism.

### The zf-UBP domain of USP22 engages in SLiM-based interaction for complex assembly

To conclude our survey, we screened the remaining 10 zf-UBP domains found in USPs for peptide binding. Most of these did not enrich any binding peptides. However, we found a small but interesting set of peptide ligands for the zf-UBP domain of USP22 (Fig. 6). USP22 is an oncogene promoting cancer progression^44,45^ and acts on histone substrates as part of the multiprotein transcriptional co-activator SAGA (Spt-Ada-Gcn5 acetyltransferase) complex, as well as on non-histone substrates^46–48^. Five potential peptide ligands were found for the USP22 zf-UBP domain of which a peptide from the Ataxin-7-like protein 1 (ATXN7L1; 36-VPSPEAFLGKPWSSWI-51) (Fig. 6A) appeared particularly relevant as it is part of the deubiquitylation module of the SAGA complex, and the USP22 zf-UBP domain is known to be crucial for interactions within the SAGA complex^48–50^. Alanine-scanning peptide array analysis of the ATXN7L1 peptide confirmed a key role of the tryptophans and revealed the contribution of upstream and downstream residues (FLGxP**W**xx**W**) (Fig. 6A). The USP22 zf-UBP-binding ATXN7L1_36-51_ complex was modelled by AF3 with high confidence (ipTM: 0.81). An extended version of the ATXN7L1 peptide (36-<u>VPS**PEAFLGKPWSSWI**</u>**DA**AKLH-57) was modelled with even higher confidence (pLDDT> 90; ipTM: 0.84) (Fig. 6B). The model suggests that the two tryptophans (W47, W50) dock into a hydrophobic cavity located on the opposite side of the typical zf-UBP binding site for the di-Gly motif of ubiquitin (Fig. 6B). The affinity of the USP22 UBP-ATXN7L1 interaction was found to be 0.8 μM, as determined by FP-monitored affinity measurements (Fig. 6C). The interaction between GST-USP22 zf-UBP and FLAG-ATXN7L1 was confirmed through GST-pulldown assays and the interaction was abolished by mutations of the binding motif (W47A/S48G/S49G/W50A) (Fig. 6D). The paralogous proteins ATXN7 and ATXN7L2, which are also part of the deubiquitylation module of the SAGA complex and are thought to be interchangeable, contain peptide regions with similar conserved GxxWxxW sequences (Fig. 6E, F). Additionally, the N-terminal region of ATXN7 has been shown to be necessary for incorporation into the DUB module of the SAGA complex^51^. The ATXN7 and ATXN7L2 peptides were confirmed to bind USP22 zf-UBP, although with lower affinity as compared to the ATXN7L1 peptide (ATXN7 K_D_≈ 11 μM, ATXN7L2 K_D_≈ 130 μM) (Fig. 6C). Finally, we note that the zf-UBP domain of the USP22 yeast orthologue Ubp8, interacts with the N-terminal region (residues 1-40) of the yeast orthologue of ATXN7/L1/L2 (SAGA associated Factor, 73 kDa (Sgf73)^52^. The interaction involves a 30-DS**W**KS**L**M-36 stretch, which corresponds to 45-KP**W**S**SW**I-51 in human ATXN7L1^53^. Thus, we conclude that the zf-UBP domain can use distinct binding sites for substrate recognition and complex assembly.

**Figure 6.**
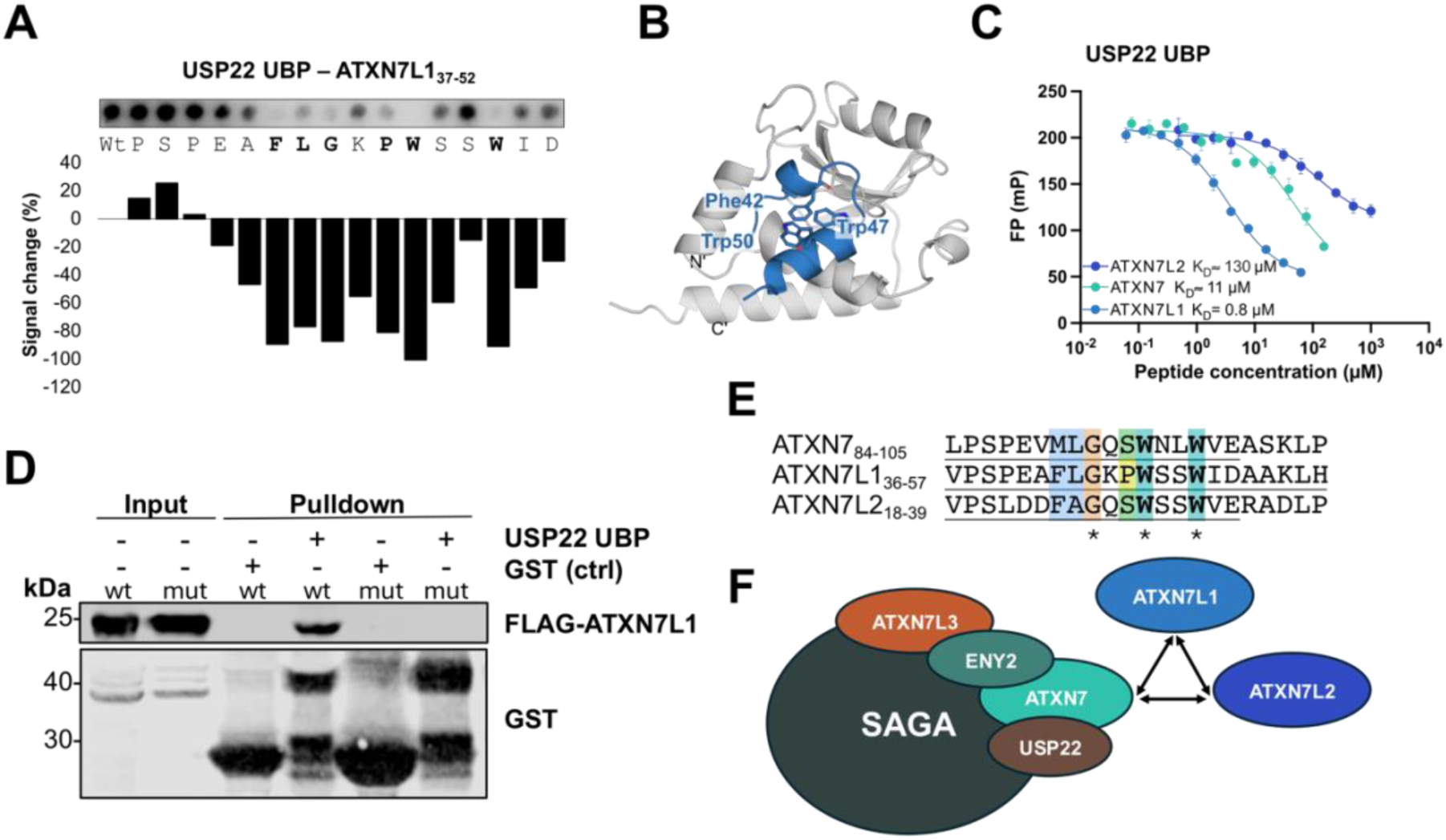
SLiM-based interactions of the USP22 zf-UBP domain. **A.** Peptide array alanine scanning of the USP22 zf-UBP binding peptide from ATXN7L1_37-52_. **B.** AF3 model of the complex between the USP22 zf-UBP domain and the ATXN7L1 peptide (ipTM: 0.8). The peptide residues are shown in stick representation. **C.** FP-monitored affinity measurements between USP22 zf-UBP domain and peptides from ATXN7_84-105_, ATXN7L1_36-57_, and ATXN7L2_18-39_. (n = 3). **D.** GST-pulldown validation of the interaction between GST-tagged USP22 zf-UBP domain and FLAG-tagged ATXN7L1 wild-type or motif mutant (W47A/S48G/S49G/W50A) from HEK293T cell lysate. **E.** Sequence alignment of the USP22 zf-UBP binding ATXN7/7L1/L2 peptides (asterisk indicates full conservation). **F.** Schematic representation of the SAGA complex, in which the ATXN7/7L1/L2 proteins are interchangeable. Schematic inspired by Felicio *et al*, 2023 ^54^.

### Concluding remarks

In this study, we systematically investigated SLiM-based interactions involving two full-length USPs and 29 ADs from the USP family of deubiquitinases. We uncovered a diverse array of binding motifs and provide novel insights into the peptide-binding capabilities of DUB ADs. Our results suggest that SLiM-binding is a relatively common function among these domains, resembling the recognition of degrons by E3 ligases.

For certain ADs, such as the zf-UBP domain of USP22, we identified only a few ligands, yet these ligands appear biologically relevant in the context of the assembly of the SAGA complex^51^. In contrast, other domains, including the zf-UBP domains of USP20 and USP33, interact with a large number of binding motifs across the proteome. This observation is consistent with their roles as broad-specificity DUBs that act on numerous substrates^30–32^. Furthermore, we show that both CYLD and USP20/33 harbour more than one SLiM-binding domain. This suggests that efficient complex assembly and substrate targeting may rely on a multicomponent recognition strategy, involving SLiM-based interactions mediated by ADs in addition to polyubiquitin chain recognition by the catalytic domain. A major challenge in establishing the functional relevance of these identified motif-DUB interactions is that the formation of the DUB-ubiquitinated substrate has to be triggered. However, for many DUBs, this information is lacking. We propose that coincidence detection, i.e., the binding of the SLiM motif by the AD plus ubiquitin on the motif-containing protein, enables the specific recognition of ubiquitinated substrates. Hence, the DUB-SLiM interaction identified in this study can guide experiments to define the cellular pathways regulated by these DUBs.

Another intriguing finding is the overlap between several identified USP20/USP33 zf-UBP binding sites and known sites of ubiquitination. This suggests multiple possibilities for the functional interplay between ubiquitin "writers" (E3 ligases) and "erasers" (DUBs). One plausible scenario is that motif binding to zf-UBP domains shields specific lysine residues from ubiquitination, thereby modulating substrate modification. Alternatively, these interactions may reflect a mechanism of processive deubiquitination. While these scenarios remain hypothetical, they point to sophisticated regulatory strategies that warrant further experimental investigation. Interestingly, Marcello Clerici and colleagues^55^ demonstrated that the DUSP domain of USP4 facilitates ubiquitin release following hydrolysis, enhancing catalytic efficiency. Furthermore, the USP4 DUSP domain works synergistically with the UBL domain to retain the ubiquitylated substrate in the active site, preventing competition from other substrates. By analogy, we hypothesise that the DUSP2 domain of USP20 may similarly collaborate with the UBP domain to confer substrate specificity towards HIF1A. HIF1A was previously identified as an interactor of USP20, and our work here provides a molecular explanation for how this interaction occurs ^42^. Other modes of regulation are of course possible, including phosphorylation sites overlapping with the binding motifs or located in the flanking regions, which may indicate further regulatory control.

In summary, our study uncovered hitherto unknown peptide binding functions for several DUBs and broadens the motif-based interactome of the DUBs. The results suggest intricate recognition mechanisms which may enhance the specificity and complex assembly of these enzymes. From a more applied perspective, there is growing interest in developing DUBTAC approaches to induce targeted protein stabilisation. However, a major challenge in this field is targeting DUBs to substrates such that the DUB is optimally positioned for efficient deubiquitylation. The reported peptides present an attractive strategy to enable DUB-target proximity that utilises the ADs and enhances selectivity while avoiding off-target effects caused by targeting via the catalytic domains.

## Materials and Methods

### Protein expression and purification

Proteins were expressed in *Escherichia coli (E.coli)* BL21-Gold (DE3) bacteria. The cDNAs encoding N-terminal 6-histidine-glutathione S-transferase tagged (6xHis-GST), 6xHis-tagged or GST-tagged protein domains in pETM33, pETM11, pGEX6P1 expression vectors respectively, were obtained from Genescript or kindly provided by MRC-PPU (Suppl. Table 1) The bacteria were transformed with the plasmid and grew at 37°C in 2YT (5 g/L NaCl, 16 g/L tryptone and 10 g/L yeast extract) supplemented with Kanamycin (30 μg/mL) until reaching an OD_600_ of 0.7. The protein expression was induced with 0.3mM isopropyl β-D-1 thiogalactopyranoside (IPTG) for 18 hours at 18°C shaking at 220 rpm. For zinc-finger domains, 50μM of ZnSO_4_ was added during expression. The bacterial cultures were pelleted for 15 minutes at 4000×*g* at 4°C and stored at −20°C until purification. The pellets were resuspended in lysis buffer (1x phosphate buffered saline (PBS), 0.5 % Triton-X, 5 mM MgCl2, 10μg/mL lysozyme, cOmplete^TM^ EDTA-free protease inhibitor cocktail (Roche), 10 μg/mL DNAseI) and incubated for 1 hour shaking at 4°C. The suspension was sonicated prior to phage selections or lysed using a cell disruptor (Constant systems) prior to affinity measurements and SPOT arrays, followed by centrifugation at 16,000×*g* for 1 hour at 4°C. The supernatant was incubated with glutathione (GSH) sepharose (Cytiva) or Ni2+ sepharose (Cytiva) for 1 hour at 4°C while shaking. The beads with the bound protein were washed with 1x PBS and eluted with 10 mM reduced glutathione in 50 mM Tris pH 8.0. For fluorescence polarization experiments the proteins were not eluted from the beads and the 6xHis-GST tag was cleaved overnight on GSH sepharose beads at 4°C while shaking with PreScission Protease in cleavage buffer (50mM Tris pH 7.5, 150mM NaCl, 1mM DTT). The sample containing the cleaved domain was separated from the tag and protease and buffer-exchanged to 50 mM sodium phosphate buffer pH 7.4 with 1mM DTT using a PD-10 desalting column (Cytiva). The purity and size of the proteins were analyzed by SDS-PAGE electrophoresis.

### ProP-PD selections

The human disorderome library was used for four rounds of phage display selections ^16^. GST-tagged, 6xHis-tagged or GST-6xHis purified proteins were used as baits and GST was used as control by immobilization on a 96-well MaxiSorp plate (Nunc) overnight at 4°C while shaking (10 μg of protein in 100 μL PBS). The wells containing the immobilized proteins were blocked with 0.5% BSA in PBS for 1 hour at 4°C while shaking. The phage library was prepared by precipitation in 1/5th volume PEG/NaCl (20% PEG-8000 + 0.4M NaCl), incubated for 10 minutes on ice followed by centrifugation at 10,000 × g for 10 minutes at 4°C and resuspended in PBS. The GST-coated wells were washed 4 times with 0.05% Tween 20 in PBS and 100 μL/well phage library was added and incubated 1hour at 4°C while shaking. The protein-coated wells were washed 4 times with 0.05% Tween 20 in PBS and the phage solution from the GST-coated wells was transferred to the protein-coated wells and incubated 2 hours at 4°C while shaking. The unbound phages were removed by washing 5 times with 0.05% Tween 20 in PBS and the bound phages were eluted with 100 μL/well log-phase E.coli OmniMAX by incubating for 30 minutes at 37°C while shaking. M13KO7 helper phages (10^11^ PFU/mL) were added to each well and incubated for 45 minutes at 37°C while shaking. The hyper-infected bacterial cultures were transferred to 1 mL 2YT supplemented with carbenicillin (100 μg/mL), kanamycin (30 μg/mL) and 0.3mM IPTG and grew overnight at 37°C while shaking. The bacterial cultures were pelleted by centrifugation at 2000 × g for 10 minutes at 4°C. The pH of the phage supernatant was adjusted with 1/10th volume of 10× PBS and inactivated any remaining bacteria by heating at 65°C for 10 minutes. The phage pools from each round were used for the next round of selections and the process was repeated four times. To evaluate the enrichment of each round of selection phage pool ELISA was performed. As before 10 μg of GST and protein were immobilized on 96-well MaxiSorp plate (Nunc) overnight at 4°C. The wells were blocked with 0.5% BSA in PBS for 1 hour at 4°C while shaking. The phage pools from each round were added to the respective GST-coated and protein-coated wells and incubated for 1 hour at 4°C while shaking. The wells were washed four times with 0.05% Tween 20 in PBS and 100 μL (1:5000) anti-M13 HRP-conjugated antibody (Nordic Biosite) was added to each well and incubated for 1 hour at 4°C while shaking. The wells were washed four times with 0.05% Tween 20 in PBS and 1 time with PBS and 100 μL TMB substrate was added to each well until a cyan color was developed. The reaction was stopped by addition of 100 μL 0.6 M H_2_SO_4_ to each well. The absorbance was measured for both GST-coated and protein-coated wells at 450 nm using the SpectraMax iD5 Multi-Mode Microplate Reader (Molecular Devices) and the binding enrichment was evaluated. The peptide-coding regions of binding-enriched phage pools were PCR-amplified and barcoded with Phusion High-Fidelity polymerase (Thermo Scientific). The PCR products were normalized with Mag-bind Total Pure NGS and cleaned-up from 2% agarose gel using the QIAquick Gel extraction Kit (Qiagen). The samples were analyzed using the Illumina MiSeq platform.

### SPOT arrays

Peptides were synthesized on cellulose membranes with standard Fluoroenylmethylox-ycarbonyl (Fmoc) chemistry using Multipep automated synthesizer at INTAVIS (Tübingen, Germany) or ordered from JPT (PepSpots). The membranes were activated with methanol for 5 minutes and washed three times for 5 minutes with TBST (50 mM Tris, 137 mM NaCl, 2.7 mM KCl, pH 8.0, 0.05% Tween-20). The membranes were incubated with blocking buffer (5% skim milk powder in TBST) for 2h at room temperature while shaking. The membranes were incubated with the GST-tagged protein diluted at the desired concentration in blocking buffer and incubated overnight at 4°C while shaking. The membrane was rinsed 3 times with TBST and incubated with HRP-conjugated anti-GST antibody (Cytiva, RPN1236; 1:3000 dilution) in blocking buffer for 1 hour at 4°C while shaking. The membrane was rinsed 3 times with TBST and chemiluminescence detection was performed using ECL reagent (Clarity Max Western ECL substrate, 1705062, Bio-Rad) and ChemiDoc Imaging system (Bio-Rad). The images were analyzed in Fiji (ImageJ2 version 2.9.0).

### Site-directed mutagenesis

The UBP domain mutations (USP20 UBP: E55K-D59R, USP33 UBP: E86K-D90R) were introduced by 2-step PCR with Pfu DNA polymerase (Agilent). After each step, the PCR product was digested with DpnI restriction enzyme (Thermo Scientific) and transformed into *E. coli*. The DNA was extracted from bacteria with QIAprep Spin Miniprep Kit (Qiagen) and the mutated sequences were confirmed by Sanger sequencing. Mutagenesis of mammalian USP20 and USP33 constructs was performed and obtained by Genescript and for mammalian HIF1A, CEP192 and APBA2 constructs was performed and obtained by MRC PPU Reagents and Services.

### Fluorescence polarization

Affinity measurements were carried out using the SpectraMax iD5 Multi-Mode Microplate Reader (Molecular Devices) at 485 nm excitation and 535 nm emission. The measurements were performed at room temperature in a non-binding black half area 96-well plate (Corning) with a total volume of 50 μL per well. Peptides were obtained from GeneCust (France) with >95% purity. FITC-labeled peptides were dissolved in dimethyl sulfoxide (DMSO) and unlabeled peptides were dissolved in 50 mM phosphate buffer pH 7.4. Saturation measurements were performed using a serial dilution of the protein domains in phosphate buffer (25 μL) and the addition of 10 nM FITC-labeled peptide (in phosphate buffer supplemented with 2mM DTT) (25 μL). Displacement experiments were performed using a serial dilution of the unlabeled peptide in phosphate buffer (25 μL) and the addition of a master mix (25 μL) containing the protein (at a concentration of two-fold the K_D_ of the FITC-labeled peptide) in complex with 10 nM FITC-labeled peptide. All measurements were performed in three technical replicates. The data were analyzed using GraphPad Prism version 9.2.0 for MacOS (GraphPad Software, San Diego, California USA, www.graphpad.com). The saturation curves were fitted to the quadratic equation, and the displacement curves were fitted to a sigmoidal dose-response equation as described previously ^56^.

### Computational modeling

Peptides binding to the zf-UBP was modeled with AF2-multimer-v3 ^57^. The DUSP2 domain was modelled with AlphaFold 3 ^17^. Interactions were analyzed with the PLIP webserver and Chimerax v1.8 was used for visualization ^58,59^. The USP25 CTD, CYLD CAP-Gly1/2/3, USP22 zf-UBP, USP19 and USP11 UBL domains were modeled using the AlphaFold 3 webserver and PyMOL was used for visualization.

### GST-Pull-down assay

HEK293T were cultured in Dulbecco’s modified Eagle medium, high glucose, GlutaMAX™ Supplement (Gibco) supplemented with 10% fetal bovine serum (Gibco) and 1% penicillin/streptomycin at 37% and 5% CO2. The cells were transiently transfected with ATXN7L1 wild-type and mutant plasmid using jetOPTIMUS® transfection reagent (PolyPlus) following the manufacturer’s instructions, the medium was changed after 16h and the cells were harvested after 48h with trypsinization. The cells were lysed for 1h rotating at 4°C with lysis buffer (50 mM NaCl, 50 mM Tris, 1 mM EDTA pH 7.4, 0.1% NP-40 (Igepal), 1 mM DTT, cOmpleteTM EDTA-free protease inhibitor cocktail (Roche), PhosSTOP™ phosphatase inhibitors (Roche), 10 μg/mL DNAseI and RNAse). The samples were centrifuged for 1h at 16,000 g and 20 μL of supernatant was added to 1-20 μL of GST- or GST-protein bound glutathione beads purified as described above. The samples were incubated overnight rotating at 4°C. The supernatant was removed after centrifugation at 3,000 rpm for 3 min and the samples were washed three times with 1 mL of wash buffer (150 mM NaCl, 50 mM Tris pH 7.4, 0.05 % NP-40 (Igepal), 5% glycerol, 1 mM DTT). Samples from input and pull-downs were mixed with 4x Laemmli Loading dye (Bio-rad) or 2x loading dye (NuPAGE), boiled at 95°C for 5 min and stored at −20°C. Western blots were performed after running the samples in SDS-PAGE gels with Kaleidoscope (Bio-rad) or PageRuler™ Prestained Protein Ladder (Thermo Scientific) and transferred to a nitrocellulose membrane using the Trans-Blot Turbo transfer system (Bio-rad). The membranes were incubated in blocking buffer (5% skim milk in PBST (1x PBS and 0.1% Tween-20). For immunoblotting, the membranes were incubated with 1:5000 primary mouse ANTI-FLAG® M2 antibody (Sigma Aldrich) overnight and 1:5000 rabbit ANTI-GST Tag antibody (Sigma Aldrich) for 1h in 2.5% skim milk in PBST. The membranes were washed 3 times for 5 min with PBST. For visualization, the membranes were incubated with 1:5000 secondary antibody Goat anti-mouse IRDye® 680RD or Goat anti-rabbit IRDye® 800CW (LI-COR) accordingly in 5% skim milk in PBST for 1 h at room temperature. The membranes were washed 3 times for 5 min with PBST and visualized by scanning at 700 nm and 800 nm with Odyssey XF (LI-COR) imaging system.

### Transient transfection for IP and IP-MS experiments

For transfection, 1×10^6^ HEK293 cells were seeded in 10 cm dishes with DMEM, 10 % FBS, 1 % L-G (without antibiotic) medium, incubated at 37 °C, with 5 % CO_2_ for 24 h. Cells were transfected with a transfection mix containing 970 mL of Opti-MEM (Thermo Fisher Scientific), 20 μL of PEI stock (1 mg/mL) and 5 mg of the desired cDNA. For co-transfection, 2.5 μg of DNA of each plasmid was used. The mix was vortexed for 15 seconds and incubated at room temperature for 15-20 minutes. The transfection mix was added to the dish and gently mixed. Plates were incubated at 37°C for 36-48 hours before harvesting. For USP33 constructs, 5 μM Bortezomib (proteasome inhibitor) was added 4 hours before harvesting the cells. To detect endogenous and transfected HIF1A, 100 μM of CoCl_2_ was added to cells overnight to mimic hypoxia, and 5 μM Bortezomib was added 4 hours before the cells were harvested.

### Immunoprecipitation of FLAG fusion proteins

FLAG fusion proteins were immunoprecipitated from HEK293 cells transfected with plasmids encoding the protein of interest using the anti-FLAG M2 affinity gel from Sigma-Aldrich. Cell pellets were lysed with 100-200 μL/per dish lysis buffer (Pierce RIPA buffer, 1X protease inhibitor (Roche), 50 unit/μL of Benzonase Nuclease) rotating at 4°C for 30-45 minutes. 20 μL of anti-FLAG M2 affinity gel slurry per plate were equilibrated with IP buffer (10 mM Tris pH 7.5, 137 mM NaCl, 0.1% NP-40). The cell lysate was clarified by centrifugation at 17,000 xg, at 4°C for 12 minutes and the supernatant was transferred to a clean, pre-chilled low-binding Eppendorf tube. The protein concentration was estimated with a Bradford assay, and equal amount of protein per IP (0.5-2 mg) diluted with 500-750 μL of IP buffer was loaded into an Eppendorf tube with equilibrated anti-FLAG M2 affinity gel and incubated in the rotator in the cold room for 2 hours. The IP samples were centrifuged at 6,500 x g at 4 °C for 30 seconds. The supernatant containing the unbound proteins was transferred to a clean tube. The resin was washed three times with 500 μL of TBS, centrifuged at 6,500 x g at 4°C for 30 seconds. The bound FLAG fusion protein was eluted with 100 μL of 150 ng/μL 3X FLAG peptide (Sigma) in TBS and incubated in the cold room for 1 h or overnight. The tube was centrifuged at 6,500 x g at 4°C for 30 seconds the eluted proteins were transferred to a clean tube. The eluted proteins were prepared for analysis by western blot or Mass Spectrometry.

### Co-Immunoprecipitation of HA and FLAG fusion proteins

HA-tagged protein was immunoprecipitated from HEK293 cells transfected with plasmids encoding the protein of interest using Pierce anti-HA magnetic beads (Thermo Scientific). Cells were lysed with 100-200 μL of lysis buffer (Pierce RIPA buffer, 1X protease inhibitor (Roche), 50 unit/mL of Benzonase Nuclease), rotating at 4°C for 30-45 minutes. 20 μL of magnetic beads slurry per 10 cm plate were equilibrated with IP buffer on a magnetic stand. The cell lysate was centrifuged at 17,000 x g, at 4°C for 12 minutes. The supernatant was transferred to a clean, pre-chilled, low-binding Eppendorf tube. The protein concentration was estimated with a Bradford assay, 1-2 mg of protein diluted with 300 μL of TBS was loaded on equilibrated magnetic beads and incubated on the rotator at 4°C for 2 hours. The IP tube was placed on the magnetic stand, and the supernatant was removed. The unbound proteins were transferred to a clean tube. The magnetic beads were washed two times with 300 µL IP buffer and one time with TBS on the magnetic stand. The bound HA-tagged protein was eluted with 100 µL of 2X LDS without reducing agent and incubated at 70 °C for 5 minutes. The tube was placed on the magnetic stand and the eluted proteins were transferred to a fresh tube. 10 μL of NuPAGE™ Sample Reducing Agent (10X) were added to the samples and stored at −20 °C for further analysis by western blot. 10 μg of protein and 15 µL of IP was loaded for input and IP samples for Western blotting.

### Western blotting

Western blot samples were resolved on an SDS-PAGE gel using MES buffer for proteins of ≤ 50 kDa, and MOPS for proteins of ≥ 50 kDa. The gel was rinsed with distilled water and transferred onto a nitrocellulose in 1X transfer buffer (Tris base 3.0 g/L, Glycine 14.4 g/L, 20% Methanol) at 100 V for 30-35 minutes in the cold room, using the Criterion Western Protein Blotter Transfer System (BioRad). After the transfer, Ponceau staining was performed to check the efficiency of the transfer. The membrane was washed with MilliQ water to remove the Ponceau stain and then incubated in the blocking buffer (5% BSA in 1X TBST) at room temperature for 1 hour, with shaking. The blocking solution was removed and then incubated with the corresponding primary antibody overnight with shaking at 4 °C. The next day, the primary antibody was removed, the membrane was washed with TBST three times at room temperature with shaking for 10 minutes. Next, the membrane was incubated with a secondary antibody diluted 1:5000 in the blocking buffer for 1 hour. The membrane was washed with TBS three times at room temperature, shaking for 10 minutes. After washing, the membrane was developed with Clarity Western ECL Substrate (BioRad) using the ChemiDoc Imaging System (BioRad).

### IP-Mass Spectrometry

Immunoprecipitation (IP) was carried out as described above, four replicates of each IP condition were performed. The eluted protein was processed following the Strap-micro protocol (Protifi). The IP elution containing 1-100 μg protein was 1:1 mixed with 2X lysis buffer (10% SDS, 100 mM TEAB pH 8.5). Subsequently, the sample was reduced, alkylated and denatured with 5 mM TCEP at 55°C for 15 minutes, 400 mM chloroacetamide at room temperature for 10 minutes, and ∼2.5% phosphoric acid, respectively. Samples were vortexed, mixed with 165 μL of 100 mM TEAB in 90% methanol and passed through an S-Trap micro column at 4,000 g for 30 seconds to trap the proteins. The column was washed with 150 μL of 100 mM TEAB in 90% methanol (at 4,000 g for 30 seconds) three times. The S-Trap column was centrifuged again and transferred to a clean 2 mL tube for digestion with 20 μL of digestion buffer (1 μg trypsin per 10 μg sample in 50 mM TEAB), at 37°C overnight in a thermomixer without shaking. The digested peptides were eluted with 40 mL of 50 mM TEAB in water, 40 mL of 0.2 % formic acid in water, and 40 mL of 50 % acetonitrile in water centrifuging at 4,000 x g for 1 minute each time. The three elutions were pooled in a low-binding tube, placed on dry ice until frozen, dried in a SpeedVac vacuum (Thermo Scientific) and proceeded with the LCMS analysis.

### LCMS Analysis

Eluted peptides were resuspended in water supplemented with 0.015% dodecyl maltoside and 0.1% formic acid and were injected on a Vanquish Neo UHPLC System operating in trap and elute mode coupled to an Orbitrap Astral Mass Spectrometer (Thermo Fisher Scientific). Peptides were loaded onto a PepMap Neo Trap Cartridge (Thermo Fisher Scientific #174500) and analysed on a C18 EASY-Spray HPLC Column (Thermo Fisher Scientific #ES906) with a 11.8 minute gradient from 1% to 55% Buffer B (Buffer A: 0.1% formic acid in water; Buffer B: 0.08% formic acid in 80:20 acetonitrile:water, 0.7 min at 1.8 µL/min from 1% to 4% B, 0.3 min at 1.8 µL/min from 4% to 8% B, 6.7 min at 1.8 µL/min from 8% to 22.5% B, 3.7 min at 1.8 µL/min from 22.5% to 35% B, 0.4 min at 2.5 µL/min from 35% to 55% B). Eluted peptides were analyzed using data-independent acquisition mode on the mass spectrometer.

### Data Analysis

Peptides were searched against the Uniprot Swissprot Human database (released on 02/01/2023) supplemented with the USP20 and USP33 mutants using DiaNN (v1.9.0)^61^ operating in library free mode. Data filtering and statistical analysis were carried out in Python (v3.9.0) using the packages pandas, numpy, sklearn, scipy, rpy2, Plotnine and Plotly and R (v4.4.3) using the package limma.

In short, protein groups identified with a single peptide or quantified in less than three replicates in at least 1 condition were filtered out. Protein group intensities were median normalized and missing values were imputed using a gaussian distribution centred on the median with a downshift of 1.8 and width of 0.3 (relative to the standard deviation).

Protein regulation was then assessed using LIMMA eBayes and P-values were adjusted using Benjamini Hochberg multiple hypothesis correction. Proteins were considered significantly regulated if their corrected P-value was smaller than 0.05 and their fold change was greater than 1.5 or smaller than 1/1.5.

## Author contribution

**AK:** Conceptualization, Investigation, Visualization, Data analysis, Writing – original draft. **ACP:** Investigation, Visualization, Data analysis. Writing**. FL:** Investigation. **JKV:** Investigation, Visualization, Data analysis, Involved in writing – original draft. **LS:** Investigation, Data analysis. **MV:** Investigation. **NED:** Investigation, Data analysis, Funding acquisition. **OSF:** Supervision, Funding acquisition. **RI**: Investigation.**PM:** Investigation. **YI:** Conceptualization, Writing – original draft, Investigation, Supervision, Funding acquisition. **YK:** Conceptualization, Supervision, Funding acquisition. Writing.

## Funding and acknowledgement

This work was supported by grants from the Swedish Research Council (YI: 2020-03380). AK, JV and AC were supported by funding from the European Union’s Horizon 2020 research and innovation programme under the Marie Skłodowska-Curie grant agreement No. 860517 (UBIMOTIF). Sequencing was performed by the SNP&SEQ Technology Platform in Stockholm. The facility is part of the National Genomic Infrastructure (NGI) Sweden and Science for Life Laboratory and is also supported by the Swedish Research Council and the Knut and Alice Wallenberg. YK is supported by funding from an ERC Consolidator grant (grant 101002428) and MRC grant MC_UU_00038/3. Research in the OSF lab was partially supported by the Israel Science Foundation (ISF) founded by the Israel Academy of Sciences and Humanities (grant no. 301/2021). We thank INTAVIS for support with peptide array synthesis.

## Data availability

The protein interactions will be submitted to the IMEx (http://www.imexconsortium.org) consortium through IntAct ^60^. ProP-PD results will be made available through the ProP-PD portal. MS data has been submitted to the PRIDE portal.

## SUPPLEMENTARY FIGURES

**Supplementary Figure 1.**
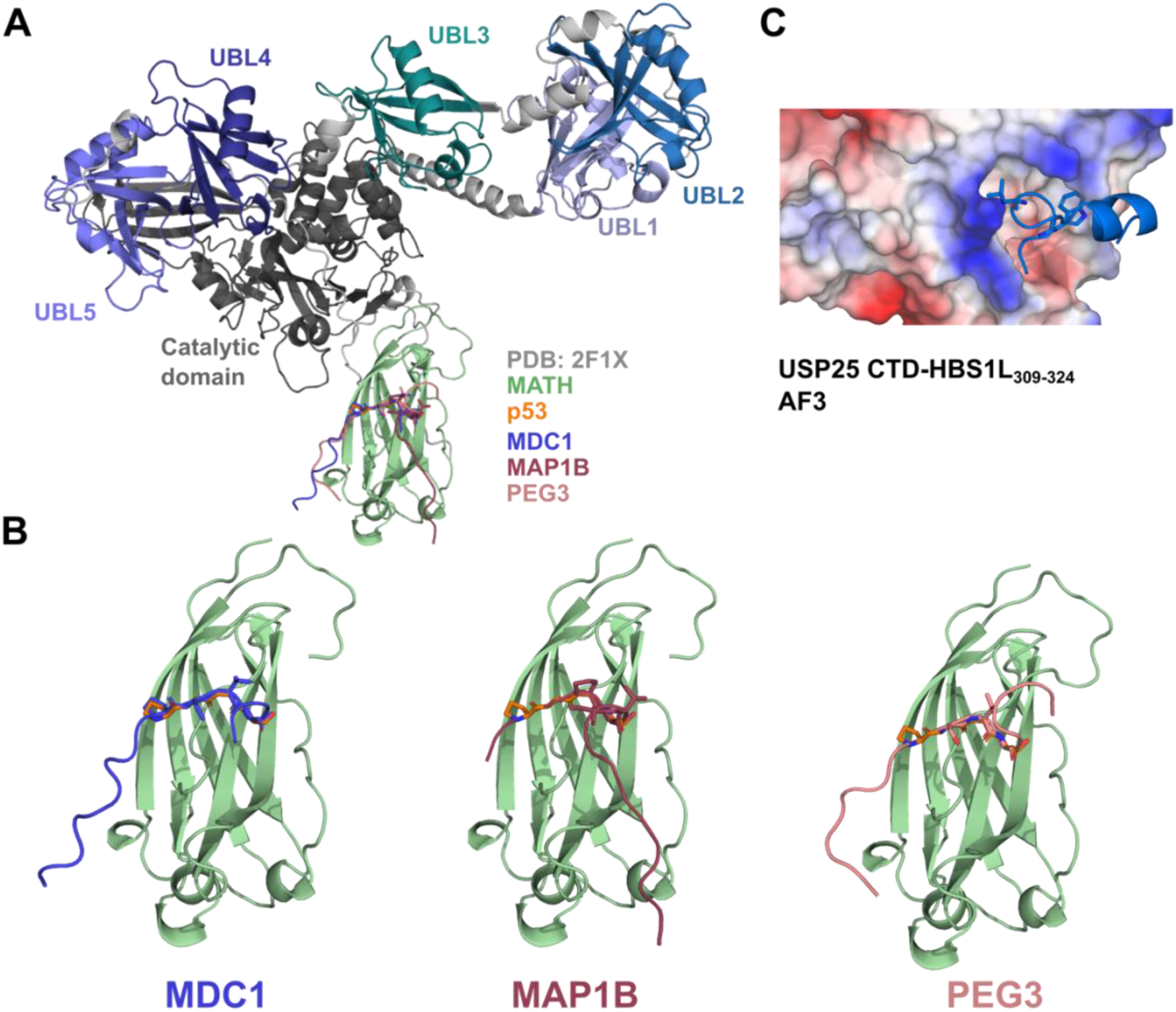
Peptides binding to the USP7 MATH and USP25 CTD domain. **A.** Crystal structure of MATH domain of USP7 bound to a p53 peptide (PDB: 2F1X) superimposed with AF3 models of the USP7 full-length predicted to bind to peptides identified in ProP-PD selections. **B.** AF3 models of the MATH domain of USP7 bound to right: MDC1, middle: MAP1B and left: PEG3 peptides. The motif key residues are shown in stick representation (see Figure 1D). **C.** AF3 model of the USP25 CTD in complex with the HBS1L_309-324_ peptide (ipTM: 0.7) with CTD domain shown with surface electrostatic potential representation.

**Supplementary Figure 2.**
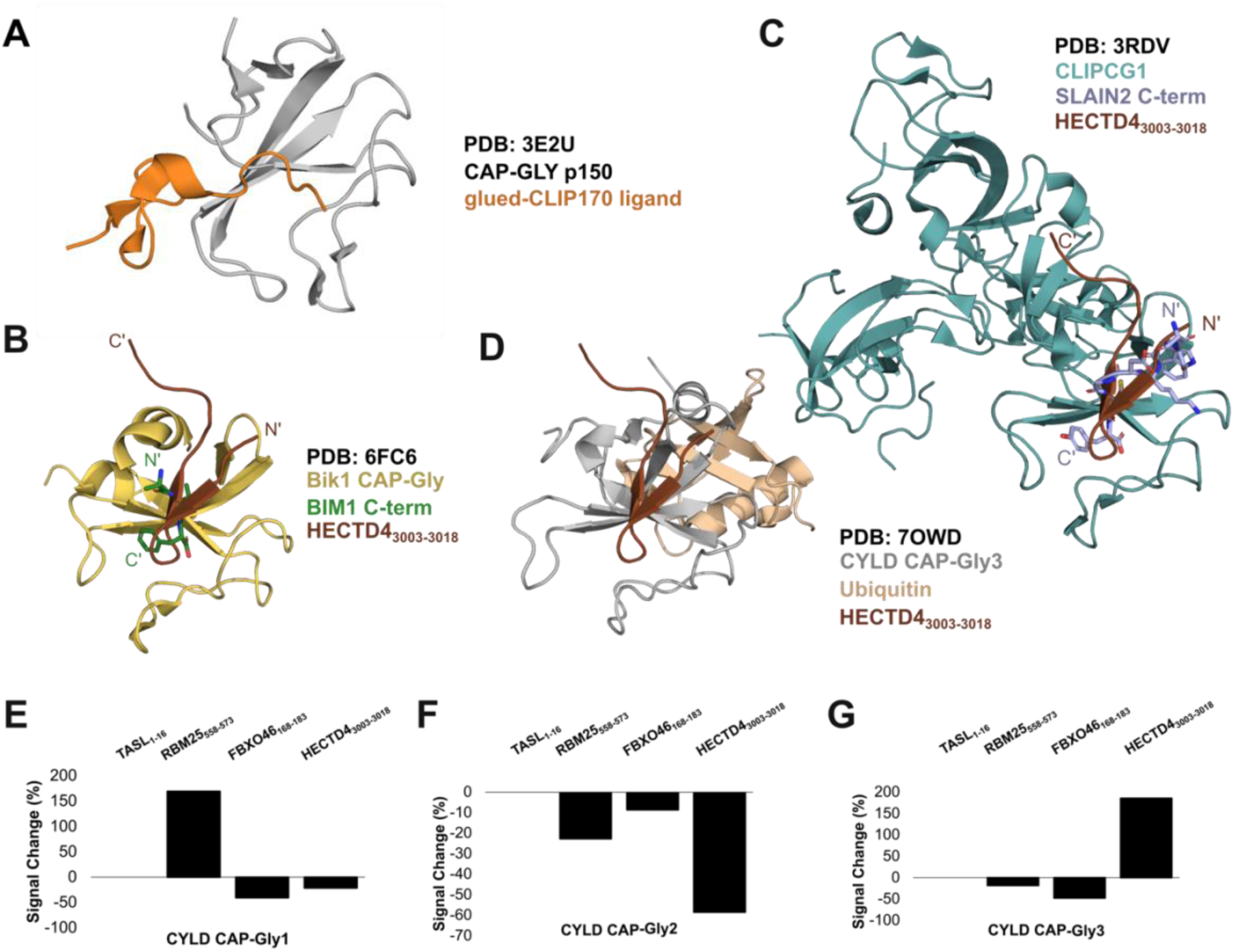
Peptide binding of CAP-Gly domains. **A.** Crystal structure of the CLIP170 peptide (orange) in complex with the CAP-Gly domain of p150 glued (PDB: 3E2U). **B.** Crystal structure of Bik1 CAP-Gly binding to BIM1 C-terminal peptide (green) (PDB: 6FC6) superimposed with the AF3 model of HECTD4_3003-3018_ peptide (brown) predicted to bind to the CAP-Gly3 domain of CYLD. **C.** Crystal structure of CLIPCG1 CAP-Gly binding to SLAIN2 C-terminal peptide (purple) (PDB: 3RDV) superimposed with the AF3 model of HECTD4_3003-3018_ peptide (brown) predicted to bind to the CAP-Gly3 domain of CYLD. **D.** Crystal structure of CYLD CAP-Gly3 binding to ubiquitin (nude) (PDB: 7OWD) superimposed with the AF3 model of HECTD4_3003-3018_ peptide (brown). **E**. CYLD CAP-Gly1 SPOT array signal intensity change (in percentage) of the wild-type TASL_1-16_ peptide compared to RBM25_558-573_, FBXO46_168-183_ and HECTD4_3003-3018_ peptides. **F.** CYLD CAP-Gly2 SPOT array signal intensity change (in percentage) of the wild-type TASL_1-16_ peptide compared to RBM25_558-573_, FBXO46_168-183_ and HECTD4_3003-3018_ peptides. **G**. CYLD CAP-Gly1 SPOT array signal intensity change (in percentage) of the wild-type TASL_1-16_ peptide compared to RBM25_558-573_, FBXO46_168-183_ and HECTD4_3003-3018_ peptides.

**Supplementary Figure 3.**
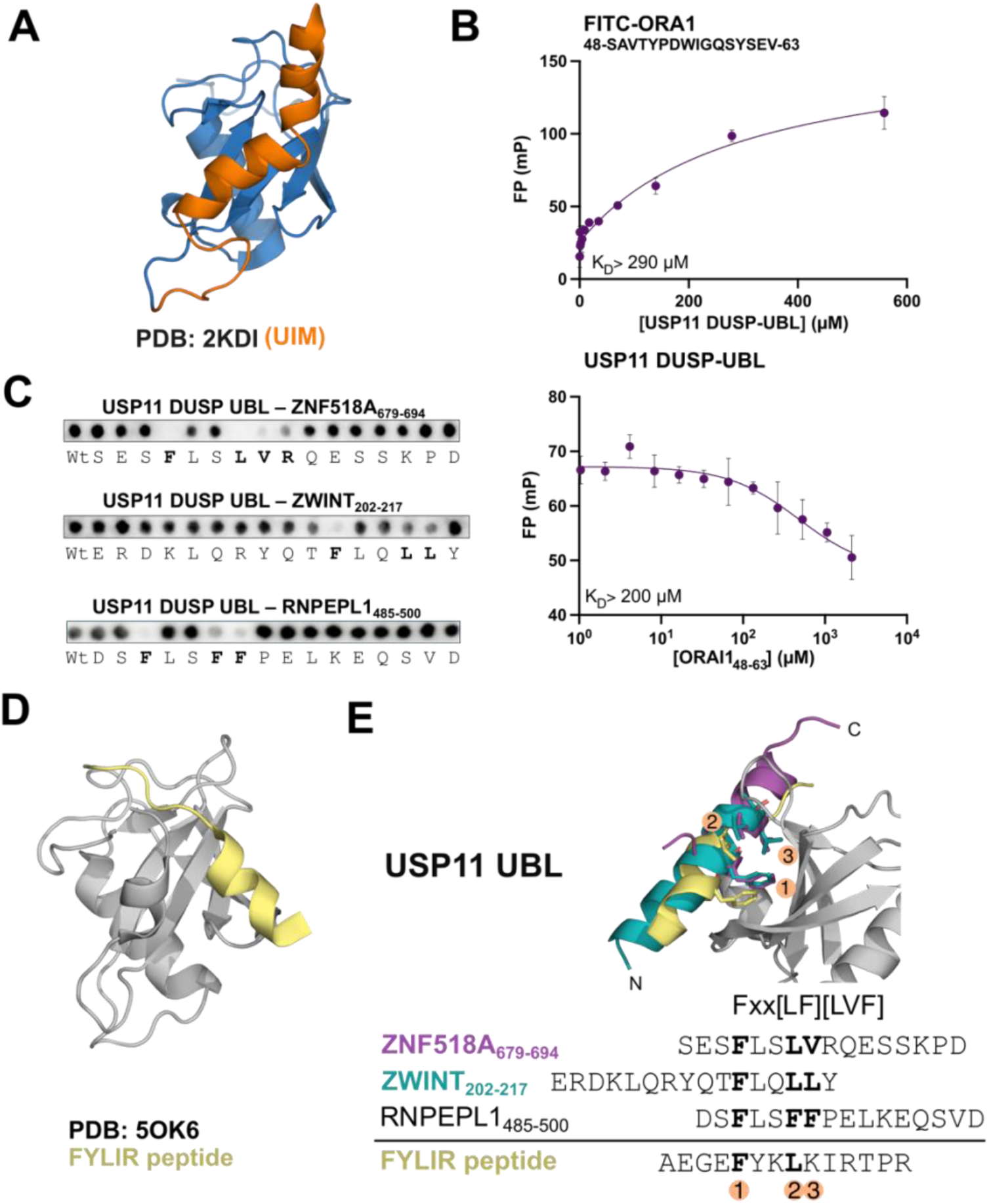
Peptide binding of the USP19 UBL and the USP11 DUSP UBL domains. **A.** NMR structure of the UIM motif binding to ubiquitin (PDB: 2KDI). **B.** FP-monitored affinity measurements of USP11 DUSP-UBL domain with labeled (direct binding) and unlabeled (Competitive binding) ORAI1 peptide (n=3). **C.** SPOT array alanine scanning of the ZNF518A_679-694_, ZWINT_202-217_, RNPEPL1_485-500_ with the USP11 DUSP UBL domain. **D.** Crystal structure of the USP11 UBL domain binding to the FYLIR peptide (PDB: 5OK6). **E.** AF3 models of the ZNF518A_679-694_ and ZWINT_202-217_ peptides superimposed with the solved structure of USP11 DUSP UBL-FYLIR complex. The Fxx[LF][LVF] motif residues are indicated and shown as sticks.

**Supplementary Figure 4.**
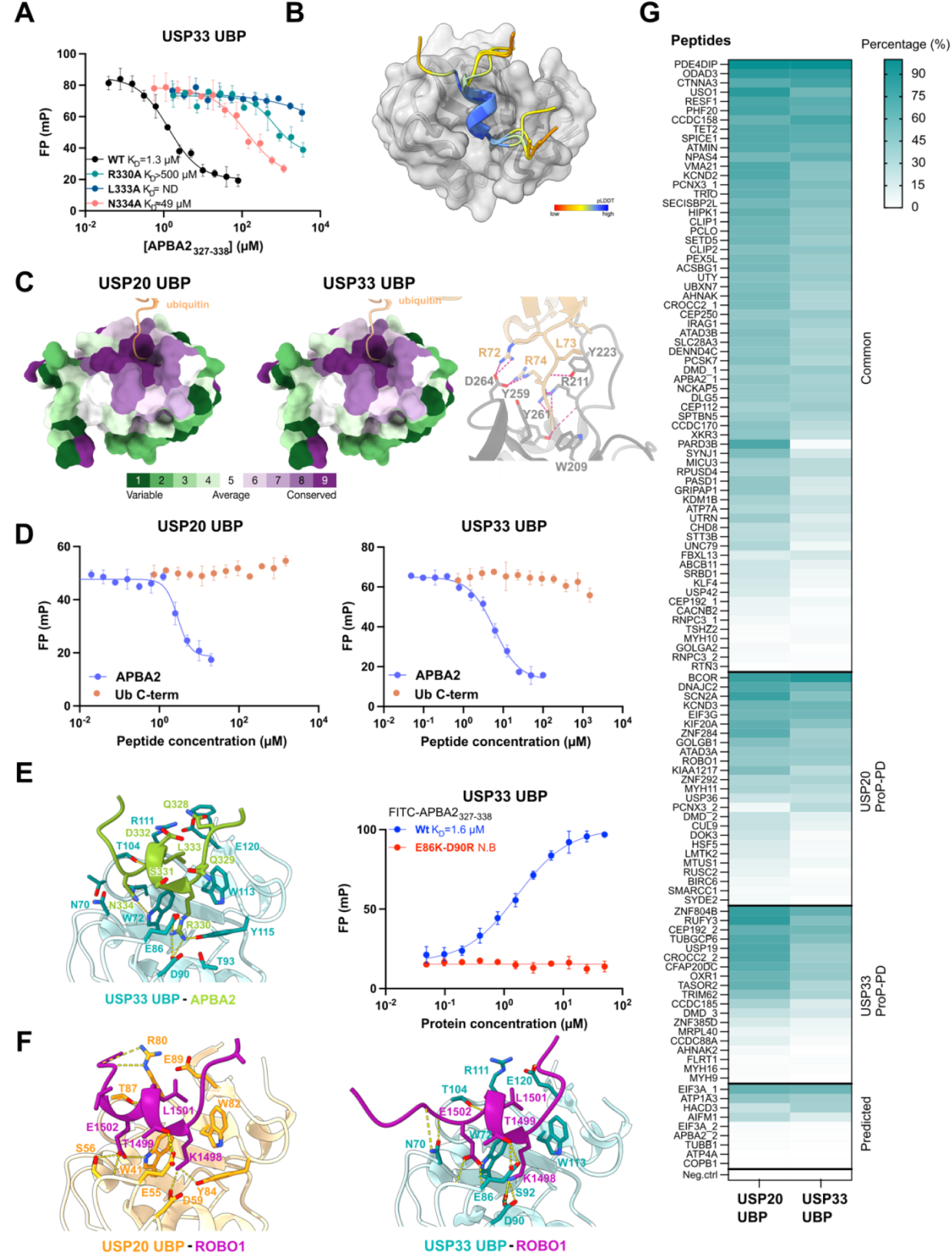
USP20 and USP33 zf-UBP domain peptide binding analysis. **A.** Competitive FP-monitored affinity measurements of USP33 zf-UBP binding to wild-type and mutant APBA2_327-338_ peptides (n = 3). ND: not determined. **B.** Superimposition of the AF3 models of the APBA2 and ROBO1 peptides in complex with USP20 and USP33 zf-UBP domains. The peptides were modeled with high confidence at the peptide region (pLDDT> 80 over the motif residues). **C.** Left, middle: Conservation of the zf-UBP domain and ubiquitin binding pocket of USP20 and USP33 zf-UBP domains. Conservation among organisms is demonstrated based on ConSurf (Yariv *et al*, 2023) Right: USP5 zf-UBP domain (grey) binding to the C-terminal tail of ubiquitin (nude) (PDB: 2G45). **D.** FP-monitored affinity measurements of the ubiquitin C-terminal peptide with USP20 and USP33 zf-UBP domains. The APBA2 peptide affinity measurements are shown for comparison (see Figure. 4H-J). **E.** Left: AF3 model of the interactions between the APBA2_327-338_ peptide and the zf-UBP domain of USP33 (pLDDT> 80 over the motif residues). Right: FP-monitored affinity measurements between wild-type and double mutant zf-UBP domains of USP33 and the APBA2_327-338_ peptide (n= 3). **F.** AF3 models of the ROBO1 peptide with the zf-UBP domains (pLDDT> 80 over the motif residues). **G.** Heatmap representation of peptide SPOT array analysis of 120 distinct peptides binding to USP20 and USP33 zf-UBP domains. Peptides were found in phage selections against both USP20 and USP33 (“Common”), in either of the two USP20 and USP33 zf-UBP domain datasets or were predicted to contain the motif. The background signal was subtracted, and the signal intensities of the peptides were normalized to the highest intensity. Peptide sequences and SPOT array results are available in the Suppl. Table 4.

**Supplementary Figure 5.**
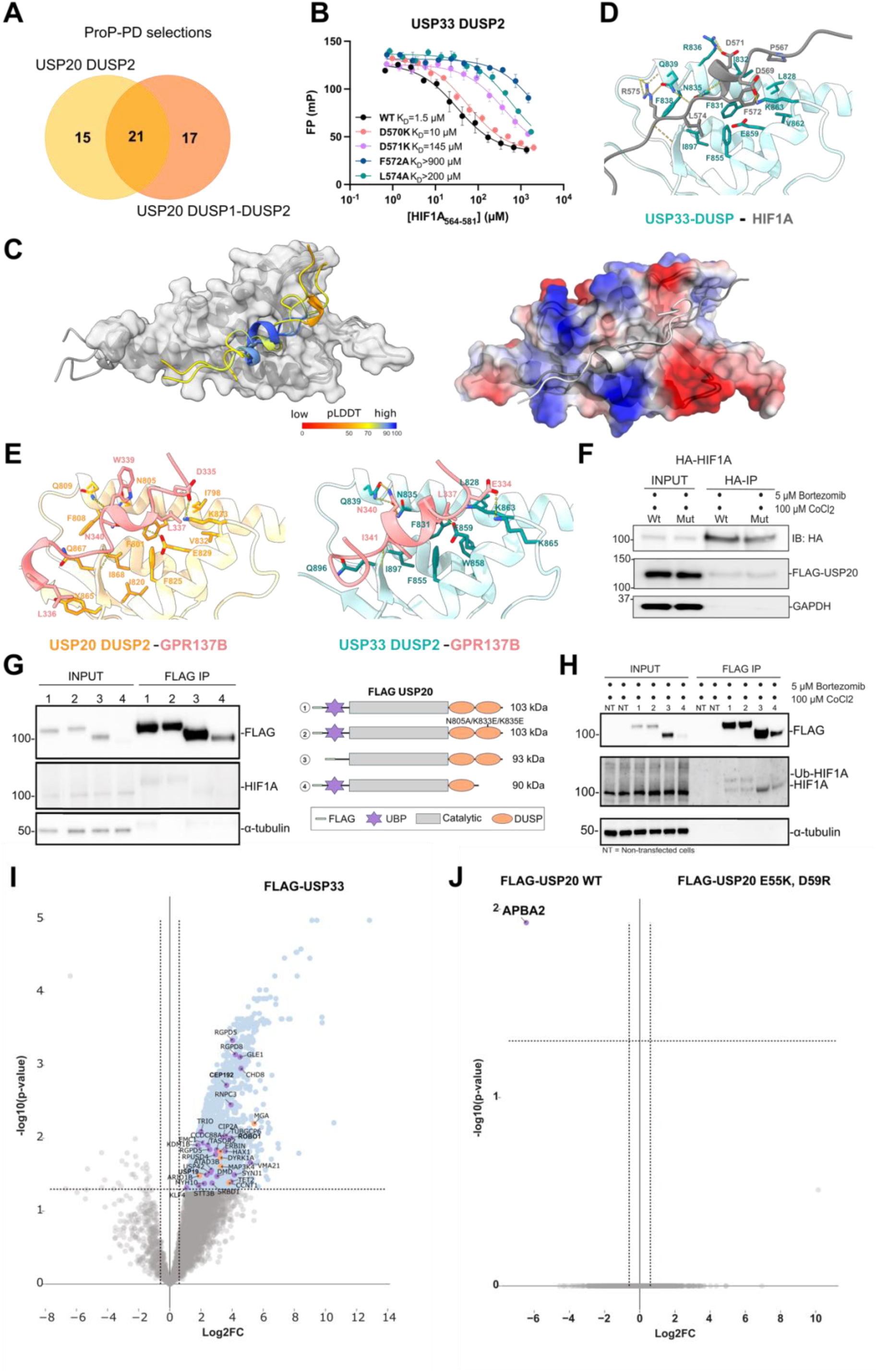
USP20 and USP33 DUSP2 domain peptide binding analysis and full-length interaction profiling. **A.** Venn diagram illustrating the overlap of peptide-binding regions identified as ligands for the USP20 DUSP2 and tandem DUSP1-DUSP2 domains through ProP-PD selections. **B.** Competitive FP-monitored affinity measurements of USP33 DUSP2 binding to wild-type and mutant HIF1A_564-579_ peptides (n = 3). **C.** Left: Superimposition of the AF3 models of the HIF1A and GPR137B peptides in complex with the DUSP2 domains. The peptides were modeled with high confidence at the [FW]x[IL] motif region (pLDDT> 90 over the motif residues). Right: AF3 model of HIF1A binding to USP20 DUSP2 in electrostatic surface potential representation. **D.** AF3 model of USP33 domain binding to HIF1A_564-579_ (pLDDT>90 values over the residues). **E.** AF3 models of the GPR137B peptide with the DUSP2 domains (pLDDT> 90 over the motif residues). **F.** Co-immunoprecipitation experiments of FLAG-USP20 with wild-type and mutant HA-HIF1A (F572A-L574A) in HEK293 cells. **G.** Left: Immunoprecipitation of endogenous HIF1A with wild-type USP20 and predicted DUSP2 binding pocket mutants or truncated zf-UBP or DUSP2 domains in normoxia. Right: Schematic representation of the USP20 FLAG-tagged constructs. **H.** Immunoprecipitation of endogenous HIF1A with wild-type USP20 and predicted DUSP2 binding pocket mutants or truncated zf-UBP or DUSP2 domains in hypoxia mimic conditions (see Suppl. Fig. 5G for descriptions of constructs used shown by numbers 1-4). **I.** FLAG-USP33 IP-MS analysis compared to the empty pCMV vector in HEK293 cells. Indicated are the interactors also identified in ProP-PD experiments for zf-UBP (purple) and DUSP2 domain (orange). **J.** IP-MS analysis of the wild-type and predicted binding pocket mutant FLAG-USP20 (E55K/D59R). The APBA2 interaction with the binding pocket mutant is significantly affected. For more information on IP-MS data (I, J) see Suppl. Table 6.

